# Astrocyte-derived PEA116 increases adult hippocampal neurogenesis and confers stress resilience

**DOI:** 10.1101/2024.09.27.615330

**Authors:** Carron Charline, Cassé Frédéric, Sultan Sébastien, Larrieu Thomas, Richetin Kévin, Espourteille Jeanne, Robadey Nicolas, Vilademunt Marta, Sagner Andreas, Petrelli Francesco, Knobloch Marlen, Beckervordersandforth Ruth, Toni Nicolas

**Affiliations:** Center for Psychiatric Neurosciences, Department of Psychiatry, Lausanne University Hospital (CHUV) and University of Lausanne, 1011 Lausanne, Switzerland; Department of Fundamental Neurosciences, University of Lausanne, 1005 Lausanne, Switzerland; Institute of Biochemistry, Friedrich-Alexander Universität, Erlangen-Nürnberg, Fahrstrasse 17, 91054 Erlangen, Germany; Department of Biomedical Sciences, University of Lausanne, 1005 Lausanne, Switzerland

## Abstract

In the dentate gyrus of the hippocampus, the neurogenic niche regulates several steps of adult neurogenesis, from the proliferation to the integration of newly formed neurons in the hippocampal network. However, the role of astrocytes in the regulation of adult neural stem cell (aNSC) proliferation is still little described. Here, we found that blocking vesicular release from astrocytes decreased cell proliferation in the dentate gyrus, resulting in impaired adult neurogenesis. Inversely, astrocyte-conditioned medium increased cell proliferation in a vesicular release-dependent manner. We identified PEA116 as a peptide released by astrocytes, that is derived from the c-terminal portion of the PEA15 protein and increased cell proliferation. PEA116 increased ERK2 phosphorylation, decreased the expression of genes involved in aNSC quiescence, resulting in aNSC quiescence exit. The ensuing increase in hippocampal neurogenesis improved resilience to chronic stress. These findings highlight a novel peptide produced by astrocytes that regulates the early steps of adult neurogenesis, with an implication for mood disorders.

## Introduction

In the mammalian brain, adult neurogenesis occurs in two distinct regions: the subventricular zone (SVZ), where neurons migrate into the olfactory bulb, and in the dentate gyrus (DG) of the hippocampus^1^. In the DG, aNSC reside in the subgranular zone (SGZ) and, upon exiting quiescence, produce transit-amplifying progenitors, with high proliferative properties. Progenitors then give rise to neuroblasts, which migrate through the granule cell layer and mature into newborn granule neurons that functionally integrate in the hippocampal network. Increasing evidence indicates that adult hippocampal neurogenesis plays a role in stress resilience: In animal models of depression, the reduced hippocampal neurogenesis is restored by antidepressant treatment^2^ and antidepressant response on mood requires intact adult neurogenesis^3,4^. Furthermore, adult neurogenesis exerts a control over the Hypothalamic-Pituitary-Adrenal (HPA) axis and blocking adult neurogenesis increases the production of corticoids upon stress and reduces the behavioral response to stress^5,6^. Inversely, an increase in neurogenesis reduces depression-like symptoms after chronic stress^7,8^. Thus, adult neurogenesis can be considered both a biomarker of brain diseases and a therapeutic target for mood disorders.

The production of new neurons is influenced by various signals produced by the direct cellular environment of the newly formed cells, the neurogenic niche, which include stem cells and progenitors, neurons, astrocytes, microglia, oligodendrocytes or blood vessels. These cells provide paracrine or juxtracrine signaling that regulate multiple aspects of adult neurogenesis, from the proliferation of aNSC to the synaptic integration of newly formed neurons^9^. One of the most important points of regulation of adult neurogenesis is the exit of quiescence. Quiescent aNSC are enriched in transcripts related to cell signaling and cell-cell communication^10^ and, owing to their complex morphology, are in contact with several cell types of the neurogenic niche, thereby enabling paracrine signaling^11,12,13^. ANSC are therefore exposed to numerous signals from the neurogenic niche, which they compute to maintain their quiescent state or resume proliferation^14^.

By their implication in neurovascular coupling, gliotransmission, blood-brain barrier integrity and neuroinflammation, astrocytes integrate niche signaling that contributes to the homeostasis of adult neurogenesis. For example, the activity of cholecystokinin interneurons stimulates adult neurogenesis through astrocytes-mediated glutamate release^15^. Furthermore, by establishing perisynaptic processes on newborn neurons^11^, astrocytes regulate the synaptic connectivity and modulate synaptic transmission^16^. In previous experiments, we found that astrocytes participate to the release of D-serine, that enables the unsilencing of immature synapses, leading to the synaptic integration and maturation of newly-formed neurons^17^. However, the role of astrocytes in the regulation of aNSC activation in the DG is still relatively poorly described.

## Results

### Astrocytes release molecules that enable adult neurogenesis

To assess the role of astrocytes-released molecules on the regulation of adult neurogenesis, we used the dnSNARE mouse model. This double-transgenic mouse results from the crossing of a GFAP-tTA mouse, in which the tet-OFF tetracycline transactivator (tTA) is expressed under the control of the glial fibrillary acidic protein (GFAP) promoter and the tetO-dnSNARE mouse, in which the dominant-negative form of the SNARE domain of synaptobrevin-2 and the reporter GFP are expressed under the control of the tet operator^18^,. dnSNARE and wild-type (WT) mice were kept under doxycycline from mating of the parental pair until 8 weeks of age, to enable brain development and postnatal neurogenesis to occur in the presence of functional astrocytes. Thirty days later, to enable complete transgene expression^17^, mice received 3 injections of the thymidine analog Bromodeoxyuridine (BrdU) (100 mg/kg, i.p.) and were perfused 24 hours later, to assess cell proliferation using immunohistochemistry. As a control, dnSNARE and WT mice received doxycycline treatment for the entire duration of the experiment (**Fig. 1A, B**.). When kept under doxycycline, WT and dnSNARE mice showed no difference in cell proliferation in the DG (**Fig. 1C**: WT: 950 ± 27.4; dnSNARE: 923.2 ± 18.7 BrdU^+^ cells/DG). However, upon doxycycline withdrawal dnSNARE mice showed a strong decrease in the total number of BrdU^+^ cells in the DG as compared to WT mice (**Fig. 1D, E**.: dnSNARE: 296 ± 20.4, WT: 707 ± 49.4 BrdU^+^ cells/DG). To further characterize the cell types in the DG that were affected by doxycycyline removal, we used immunohistochemistry. Stem cells, identified by Sox2 expression and their nucleus in the SGZ, displayed no difference in number between dnSNARE and WT mice with or without doxycycline administration (**Fig. 1F-H**. On Dox: WT: 1201.2 ± 51.6; dnSNARE: 1246.35 ± 46.5; Off dox: WT: 1075.2 ± 55.6; dnSNARE: 1082.4 ± 70 cells/DG). In contrast, the number of Tbr2^+^ progenitors displayed a marked reduction upon doxycycline withdrawal in dnSNARE mice as compared to WT mice (**Fig. 1I-K**. On Dox: WT: 751.7 ± 28.4; dnSNARE: 815 ± 26.2; Off dox: WT: 651.16 ± 21.8; dnSNARE 193.3 ± 23.1 cells/DG). Thus, ablation of astrocytic vesicular release reduced cell proliferation in the DG by reducing the number of the highly proliferative Tbr2^+^ progenitors. Next, we examined the effect of the inhibition of astrocytic-vesicular release on DCX^+^-expressing neuroblasts and immature neurons (**Fig. 1L, M**). Thirty days after doxycycline removal, dnSNARE mice displayed a non-significant decrease in the number of DCX^+^ cells as compared to WT mice (**Fig.1N**. dnSNARE: 3616.2 ± 289.4; WT: 4648 ± 373.9), which became significant 60 days after doxycycline removal (**Fig. 1O**; dnSNARE 1200 ± 76.9; WT 4100 ± 448.4 cells/DG).

**Figure. 1:**
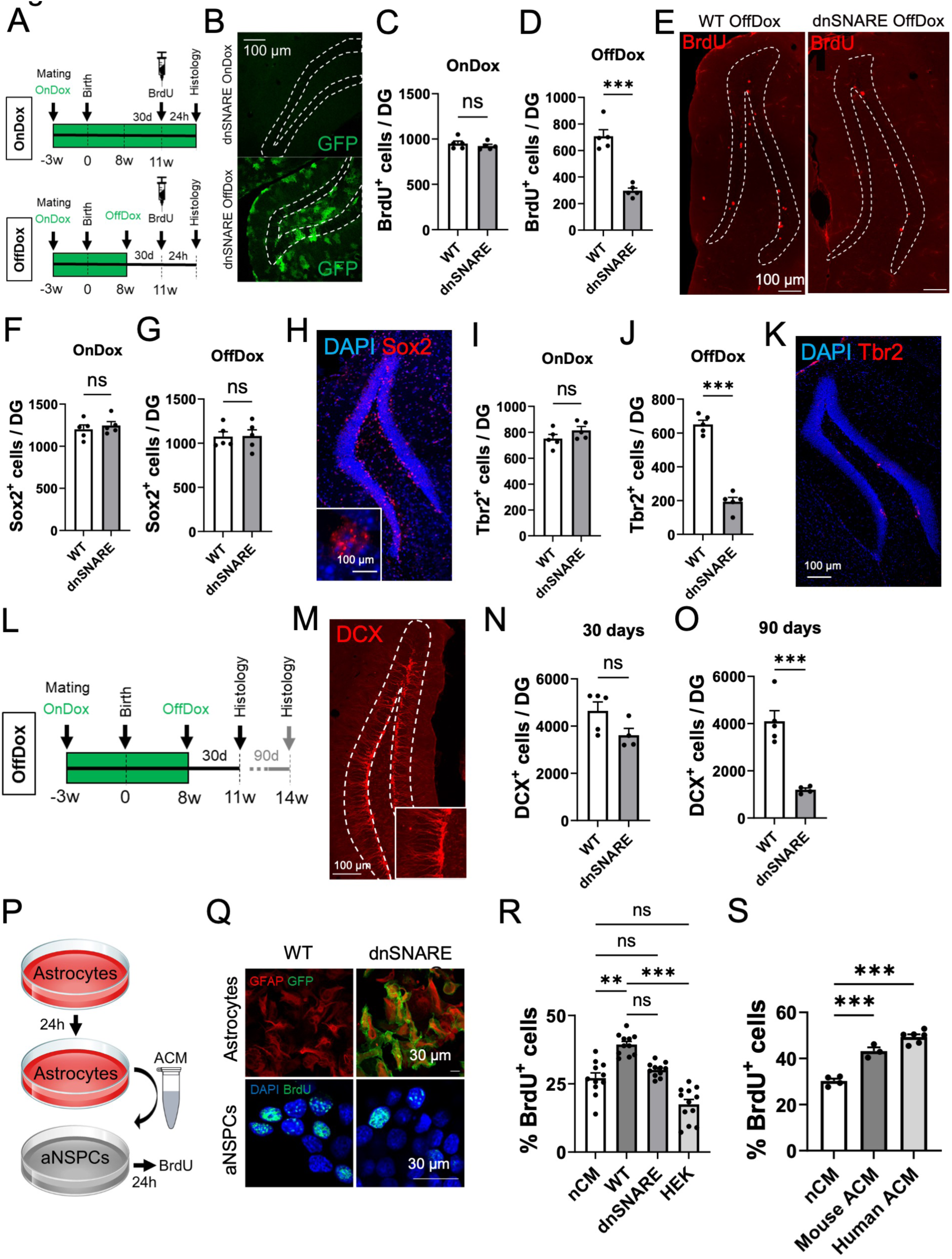
Astrocytes release soluble molecules that increase adult neurogenesis. **A.** Experimental design for the quantification of aNSPC proliferation and number in dnSNARE mice. **B**. Representative confocal micrographs of hippocampal slices stained for BrdU (red) and GFP (green) as a reporter for the expression of the dnSNARE transgene. **C.** Number of aNSPC that incorporated BrdU in the DG of WT or dnSNARE mice under doxycycline (OnDox) administration (t = 0.8071, p = 0.4429, unpaired t-test, two-tailed, n = 5 mice per group) or **D.** 30 days after after the removal of doxycycline (OffDox) (t = 7.683, p < 0.001, unpaired t-test, two-tailed, n = 5 per group). **E.** Confocal micrographs of BrdU staining in WT (left) and dnSNARE mice (right). **F-G**. Number of Sox2^+^ stem cells in the DG of WT or dnSNARE mice under doxycycline (**F.** OnDox, t = 0.6501, p = 0.5338, unpaired t-test, two-tailed, n = 5 per group), and upon doxycycline removal (**G.** OffDox, t = 0.08078, p = 0.9376, unpaired t-test, two-tailed, n = 5 per group). **H.** Confocal micrograph of Sox2 immunostaining. **I-J**. Number of Tbr2^+^ progenitor cells in the DG of WT or dnSNARE mice under doxycycline administration (**I.** OnDox, t = 1.466, p = 0.1809, unpaired t-test, two-tailed, n = 5 per group) and after doxycycline removal (**J.** OffDox, t = 12.88, p < 0.001, unpaired t-test, two-tailed, n = 5 per group). **K.** Confocal micrograph of Tbr2 staining. **L**. Experimental design for the study of neuroblasts/immature neurons number in dnSNARE mice. **M**. Confocal micrograph of DCX staining. **N-O.** Number of neuroblasts/immature neurons (DCX^+^) in the DG of WT or dnSNARE 30 days after doxycycline removal (**N.** t = 2.087, p = 0.0753, unpaired t-test, two-tailed, WT: n = 5, dnSNARE: n = 4) and 90 days after doxycycline removal (**O.** t = 5.653, p = 0.0008, unpaired t-test, two-tailed, n = 5 per group). **P.** Schematic illustration of the production of astrocytes conditioned medium (ACM). **Q.** Confocal micrographs of WT and dnSNARE astrocytes *in vitro* stained for GFAP (red) and reporter expression (GFP, green, upper panels) and of aNSPC (BrdU, green) treated with ACM from WT and dnSNARE astrocytes (lower panels). **R**. Percentage of proliferating aNSPC after exposure to non-conditioned medium (n=11 replicates), medium conditioned by WT astrocytes (n = 11 replicates), dnSNARE astrocytes (n = 11 replicates) or HEK-293 cells (n = 12 replicates) (F = 34.23, p < 0.001, non-parametric Kruskal-Wallis test followed by Dunn’s multiple comparisons test). **S**. Percentage of proliferating aNSPC after exposure to non-conditioned medium (nCM, n = 4 replicates), medium conditioned by mouse astrocytes (n = 3 replicates) or human astrocytes (n = 6 replicates) (F=65.59, p<0.001, One-Way ANOVA followed by Tukey’s multiple comparisons test). Histograms show average ± SEM. ns.: not significant p>0.05; *p<0.05; **p<0.01; ***p<0.001.

Together, these results suggest that blocking vesicular release from astrocytes reduced cell proliferation and the number of immature neurons in the DG, in line with the reduced adult neurogenesis that we previously reported in dnSNARE mice^17^. However, the GFAP promoter being active in aNSC, dnSNARE expression in aNSC may interfere with vesicular release from these cells, which is known to participate to the regulation of adult neurogenesis^9^. To assess the role of astrocytes-released molecules on aNSC proliferation, we designed an *in vitro* assay in which culture medium conditioned by purified astrocytes (ACM) was applied to adult hippocampal stem/progenitor cells (aNSPC) and their proliferation was evaluated using BrdU incorporation (**Fig. 1P, Q**). ACM from WT astrocytes significantly increased the number of proliferative aNSPC as compared to non-conditioned medium (nCM). In contrast, no increase in proliferation was observed with ACM from dnSNARE astrocytes or medium conditioned by HEK-293 cells (**Fig. 1R**. nCM: 27.05 ± 1.9, WT: 39.31 ± 1.2, dnSNARE: 30.08 ± 0.8, HEK-293: 17.4 ± 1.8 BrdU^+^ cells/DG). ACM from human astrocytes produced a similar increase in the number of proliferative aNSPC as ACM from mouse astrocytes (**Fig. 1S**. nCM: 30.2 ± 1, Mouse ACM: 43.1 ± 1.7, Human ACM: 49.3 ± 1.1 BrdU^+^ cells/DG). Thus, astrocytes release molecules that increase neural stem/progenitor cell proliferation *in vivo* and *in vitro*.

### The astrocyte-derived peptide PEA116 increases adult neurogenesis

To identify the proneurogenic molecules released by astrocytes, we used a fractionation assay. ACM was filtered using a 3 kiloDalton (kDa) molecular weight cutoff centrifuge concentrator and either fraction (<3KDA and >3kDa) was applied to aNSPC. To avoid the depletion of molecules contained in the culture medium, both fractions were complemented with the missing fraction of nCM (>3kDa or <3kDa, respectively, **Fig. 2A**) and applied to aNSPC. Both ACM and the < 3KDa fraction of ACM increased aNPSC proliferation, whereas the >3 kDa fraction of ACM decreased aNSPC proliferation as compared to nCM (**Fig. 2B**. nCM: 33.7 ± 1.3, ACM: 39.9 ± 1.4, <3kDa ACM: 43.2 ± 1.6, >3kDa ACM: 28.0 ± 1.6 BrdU^+^ cells/DG). This result suggests that the proneurogenic molecules released by astrocytes have a molecular weight smaller than 3 kDa. To assess the nature of these molecules, we treated ACM <3 kDa fraction with proteinase K (100 μg/ml for 3 h at 37°C), then re-filtered the sample with a 3kDa filter to remove proteinase K. The resulting fraction was complemented with >3 kDa fraction of nCM and applied to aNSPC for 24h (**Fig. 2C**). As expected, both ACM and <3 kDa ACM increased aNSPCs proliferation. However, when treated with proteinase K, ACM <3 kDa did not exhibit any effect on aNSPC proliferation. Similarly, nCM filtered with a 3 kDa filter and treated with proteinase K did not modify cell proliferation (**Fig. 2D**. nCM: 32.1 ± 1.4, ACM: 43.2 ± 0.9, ACM <3 kDa: 44.9 ± 1.2, ACM <3 kDa + Prot K: 31.9 ± 1.9, nCM + Prot K: 33.6 ± 1.3 BrdU^+^ cells/DG). Thus, peptides smaller than 3 kDa mediate the proliferative effect of ACM on aNSPC. To identify the peptides that increase aNSPC proliferation, we submitted astrocytes-conditioned solution (ACS, in which cell culture medium was replaced by a defined Tyrode’s solution) to untargeted mass spectrometry-assisted proteomic analysis. We conducted analysis from five distinct WT astrocyte cultures, each derived from a separate animal. Peptides identified with identical sequences, a relative abundance superior to a threshold of 1 and present in at least three experiments were synthesized. Five peptides corresponded to these criteria (**Supp. Fig. 1A**). They were then synthesized and incubated *in vitro* on aNSPCs at a concentration of 10 μM for 24h to evaluate their effect on cell proliferation. Only peptide 5 increased cell proliferation as compared to vehicle (**Fig. 2E**. Vcl: 30.9 ± 0.47, P1: 33.2 ± 0.9, P2: 33.5 ± 1.7, P3: 31.6 ± 1.1, P4: 30.8 ± 1.3, P5: 47.9 ± 1.1 BrdU^+^ cells/DG). Peptide 5 consists of 12 amino acids (SEEEIIKLAPPP). It is derived from the C-terminal part (aa 116-127) of the 15 kDa phosphoprotein enriched in astrocytes (PEA-15, 130 amino acids) and was therefore named PEA116. Interestingly, P4, which originates from the N-terminal portion of PEA-15 (aa 18-26) did not increase aNSPC proliferation (**Fig. 2E**). The amino-acid sequence of PEA116 is conserved between human and mice. Although PEA-15 is enriched in astrocytes, the protein is not limited to this cell type and is expressed across various cell types. RNA expression databases indicate a higher expression of PEA-15 in neural tissue compared to other parts of the organism in both species. To assess PEA-15 expression in the human and mouse hippocampus, we interrogated two published data sets, one of single nucleus RNA-sequencing of human hippocampal cells^19^ and one of single cell RNA-sequencing of mouse DG cells ^20^. In both human and mouse, PEA-15 mRNA is expressed in most hippocampal cells, including astrocytes (**Fig.2F-I**). The presence of PEA116 peptide in the mouse DG was further confirmed by microdialysis and displayed similar concentrations between sedentary mice and mice exposed to exercise-induced increase in neurogenesis (**Supp. Figure 1B-D**, No run: 0.5 ± 0.1; run: 0.4 ± 0.08 fmols. No run: 1151 ± 58.1; run: 1738 ± 131.8 Ki67^+^ cells/DG). To test the possibility that PEA116 may also be present in the interstitial fluid of the Human brain, we examined the composition of the extracellular fluid collected from postmortem brains using LC/MS. We found the presence of the partial sequence of PEA116 (SEEEIIK) in extracellular vesicles purified from the brain of healthy control patients. Consistent with this observation, a similar sequence of the peptide (QPSEEEIIK) was found in human CSF in a published proteomics anaysis^21^, indicating that across species, PEA15-derived peptides that present similarities with PEA116, are expressed in the brain extracellular space.

**Figure 2:**
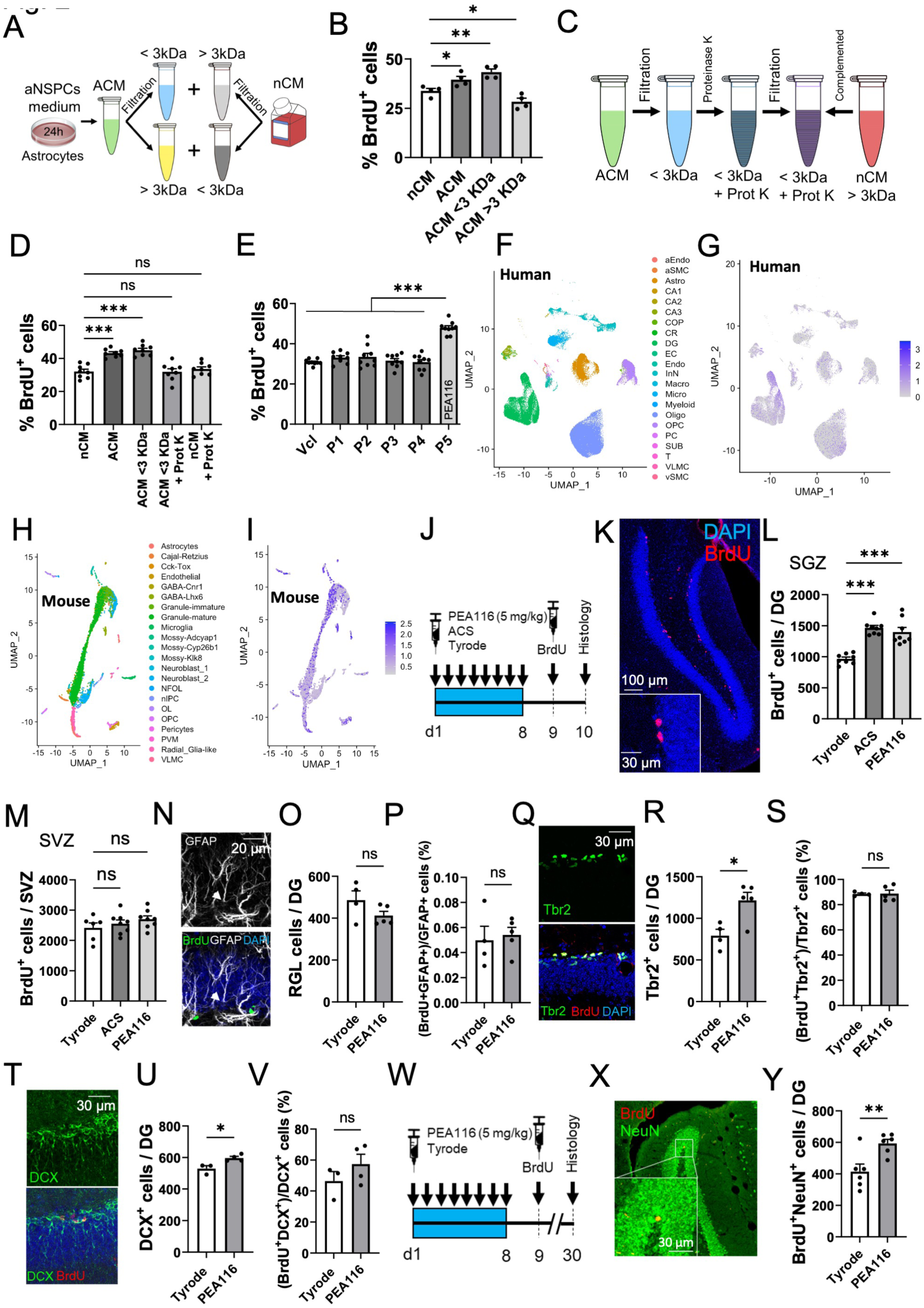
PEA116 is produced by astrocytes and increases adult neurogenesis. **A.** Schematic illustration of the mass exclusion filtering strategy for ACM. **B**. Percentage of proliferating aNSPC after exposure to non-conditioned medium (nCM), non-filtered ACM (ACM), small fraction of ACM (ACM < 3kDa) or large fraction of ACM (ACM > 3kDa) (n = 4 replicates per condition, F = 20.29, p < 0.001, One-Way ANOVA followed by Dunnett’s multiple comparisons test). **C**. Schematic illustration of the filtered ACM treated with proteinase K. **D**. Percentage of proliferating aNSPC after exposure to nCM, ACM, ACM < 3kDa, ACM< 3kDa treated with proteinase K (ACM < 3kDa + Prot K) or non-conditioned medium treated with proteinase K (nCM + Prot K) (F = 21.61, p < 0.001, One-Way ANOVA followed by Tukey’s multiple comparisons test, n = 8 wells per condition). **E**. Percentage of proliferating aNSPC after exposure to Vehicle (Vcl), P1, P2, P3, P4 or P5 peptides (F = 24.24, p = 0.0002, non-parametric Kruskal-Wallis test followed by Dunn’s multiple comparisons test, n = 9 wells per condition). **F-I.** UMAP-based dimensionality reduction of scRNAseq data of the human. **F, G.** and mouse. **H, I.** adult DG, showing cell type annotations (**F, H)** and PEA15 expression levels (**G, I). J.** Experimental design for the study of aNSPC proliferation in mice treated with vehicle (Tyrode’s solution), astrocytes conditioned solution, (ACS) or PEA116 **K.** Confocal micrograph of a hippocampal slice stained for DAPI (blue) and BrdU (red). **L**. Number of aNSPCs that incorporated BrdU in the DG of mice after treatment (F = 27.72, p < 0.001, One-Way ANOVA followed by Tukey’s multiple comparisons test, n = 8 mice per group). **M**. Number of aNSPC that incorporated BrdU in the SVZ of mice after treatment (F = 1.202, p = 0.3224, One-Way ANOVA followed by Tukey’s multiple comparisons test, Tyrode: n = 6, ACS: n = 8, PEA116: n = 8 mice). **N**. Confocal micrographs of GFAP^+^ Radial Glial-like stem cells (RGL) immunostained for BrdU. **O.** Number of RGL in the DG after treatment (t = 1.646, p = 0.1437, unpaired t-test, two-tailed, Tyrode: n = 4, PEA116: n = 5 mice). **P**. Proportion of proliferating RGL stem cells (t = 0.3529, p = 0.7346, unpaired t-test, two-tailed, Tyrode: n = 4, PEA116: n = 5 mice). **Q**. Confocal micrographs of Tbr2^+^ progenitor cells. **R.** Number of progenitor cells in the DG after treatment (t = 3.255, p = 0.014, unpaired t-test, two-tailed, Tyrode: n = 4, PEA116: n = 5 mice). **S**. Proportion of proliferating Tbr2^+^ progenitor cells (t = 0.1002, p = 0.9230, unpaired t-test, two-tailed, Tyrode: n = 4, PEA116: n = 5 mice). **T**. Confocal micrographs of DCX^+^ neuroblasts/immature neurons. **U.** Number of DCX^+^ cells in the DG after treatment (t = 3.373, p = 0.0198, unpaired t-test, two-tailed, Tyrode: n = 3, PEA116: n = 4 mice). **V.** Proportion of proliferating DCX^+^ cells (t = 1.210, p = 0.2803, unpaired t-test, two-tailed, Tyrode: n = 3, PEA116: n = 4 mice). **W**. Experimental design for the examination of net neurogenesis. **X**. Confocal micrograph of newborn neurons (BrdU^+^NeuN^+^ cells). **Y.** Number of newborn neurons in the DG of mice after treatment (t = 3.297, p = 0.0081, unpaired t-test, two-tailed, n = 6 mice per group). Histograms show average ± SEM. ns.: not significant p>0.05; *p<0.05; **p<0.01; ***p<0.001.

Next, we assessed the effect of PEA116 on adult neurogenesis *in vivo*. To this aim, 8-weeks-old mice were injected with either Tyrode’s solution, ACS or a PEA116 (5 mg/kg, i.p.), daily for 8 consecutive days. One day after the final injection, all mice received three injections of BrdU (100 mg/kg, i.p.) at 2-hour intervals and were sacrificed 24h later to assess cell proliferation in the SGZ and in the SVZ (**Fig. 2J, K**). In the DG, ACS and PEA116 produced a similar increase in proliferation as compared to Tyrode’s solution (**Fig. 2L**, Tyrode: 966.1 ± 28.9, ACS: 1465 ± 40.4, PEA116: 1399 ± 73.9 BrdU^+^ cells/DG). In the SVZ, neither ACS nor PEA116 affected proliferation, indicating that the actions of ACS and PEA116 were specific to the hippocampus (**Fig. 2M**, Tyrode: 2413 ± 165.8, ACS: 2553 ± 121.3, PEA116: 2702 ± 106.7 BrdU^+^ cells/SVZ). Next, we assessed the effect of PEA116 on different cell populations of the neurogenic lineage using immunohistochemistry. PEA116 did not modify the number of radial glia-like (RGL) stem cells, identified by a GFAP-immunopositive soma in the SGZ and a process extending radially in the granule cell layer (**Fig. 2N, O**. Tyrode: 486.3 ± 44.2; PEA116: 412 ± 20.3 RGL cells/DG). PEA116 did not modify the proliferation of RGL cells (**Fig. 2P**. Tyrode: 54.7 ± 12.5; PEA116: 59.7 ± 6.3 BrdU^+^ RGL/RGL^+^ cells (%)). However, we observed a strong increase in the number of Tbr2-expressing progenitors with PEA116 (**Fig. 2Q, R**. Tyrode: 791.9 ± 76.6; PEA116: 1215 ± 98 Tbr2^+^ cells/DG), but not in their proliferation (**Fig. 2S**. Tyrode: 88.5 ± 0.7; PEA116: 88.8 ± 2.7 BrdU^+^Tbr2^+^/Tbr2^+^ cells (%)). Similarly, PEA116 produced a small increase in the number of DCX^+^ neuroblasts and immature neurons (**Fig. 2T, U**. Tyrode: 529.5 ± 18.7; PEA116: 596.7 ± 10.4 DCX^+^ cells/DG) but not in their proliferation (**Fig. 2V**. Tyrode: 46.4 ± 6.1; PEA116: 57.4 ± 6.3 % of BrdU^+^DCX^+^/DCX^+^ cells). Finally, PEA116 increased the number of BrdU^+^NeuN^+^ newborn neurons, as observed 30 days after treatment initiation (**Fig. 2W-Y**. Tyrode: 414.4 ± 47.6; PEA116: 593 ± 25.9 BrdU^+^NeuN^+^ cells/DG). Thus, the increased neurogenesis conferred by PEA116 administration resulted mainly from an increased number of Tbr2 and DCX-expressing progenitors.

### PEA116 induces aNSC quiescence exit

The bidirectional communication between astrocytes and microglia is involved in the regulation of regulation inflammation, which impacts adult neurogenesis^22,23^. Therefore, the effect of PEA116 on neural progenitors may be indirect and mediated by a reduction in inflammation. To test this possibility, we examined the effect of PEA116 on the expression of inflammatory markers. PEA116 did not modify the number of GFAP-expressing astrocytes, Iba1-expressing microglia, the intensity of Iba1 immunostaining, the staining intensity of the inflammatory marker CD68 staining or microglia morphology and slightly decreased GFAP immunostaining intensity, suggesting that PEA116 did not modify inflammation in the DG (**Supp. Fig.2A-I**). Next, we assessed the effect of PEA116 on the fate of aNSPC *in vitro*, using live cell imaging (**Fig.3A**). Cells were imaged every 30 minutes, starting 4 hours before treatment and until 44 hours after treatment with 10μM PEA116, for a total of 48h. Cells were then treated with BrdU and fixed 1h later. We examined the fate of 60 vehicle-treated cells and 100 PEA116-treated cells that were randomly chosen among cells that remained visible within the imaging frame throughout the imaging session. We assessed cell division, cell cycle duration, the number of progenies per clone, the number of non-mitotic cells and the number of dying cells (**Fig. 3B**). PEA116 reduced the number of dying cells (**Fig. 3C**. Vehicle: 0.3 ± 0.05; PEA116: 0.09 ± 0.04 % of dead/total cells) and decreased the proportion of quiescent cells (**Fig. 3D**. Vehicle: 0.12 ± 0.03; PEA116: 0 ± 0 non mitotic/total cells (%)). PEA116 did not modify the number of progenies per cell (**Fig. 3E**. Vehicle: 2.7 ± 0.3; PEA116: 3.4 ± 0.3 clones/cell), nor the length of the cell cycle (**Fig. 3F**. Vehicle: 20.8 ± 1.8; PEA116: 19.2 ± 1.1 hours). The BrdU incorporation assay confirmed that PEA116 increased the proportion of BrdU^+^ cells (**Fig. 3G**. Vehicle: 0.33 ± 0.004; PEA116: 0.48 ± 0.01 % of total cells). These results indicate that the increase in BrdU incorporation produced by PEA116 treatment resulted mainly from increased quiescence exit and decreased cellular death of progenies.

**Figure 3:**
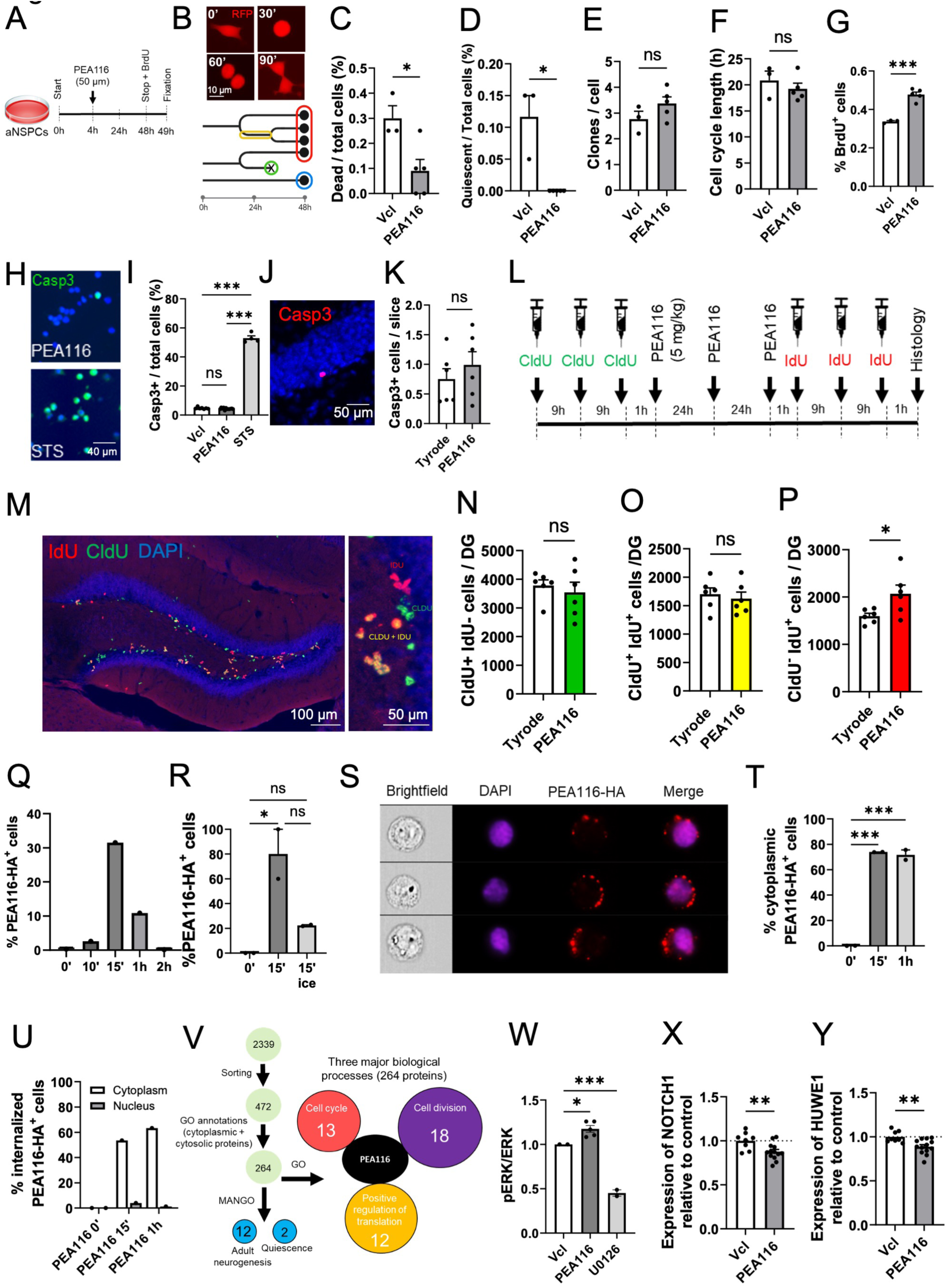
PEA116 induces aNSC quiescence exit. **A.** Schematic illustration of the live imaging of aNSPC *in vitro*. **B**. Illustration of a cell in division (top) and an example of a cell fate lineage tree (down), red = clones number, yellow = cell cycle length, green = dead cells, blue = non-mitotic cells. **C**. Proportion of cells that died among the total number of clones after the exposure to vehicle (Vcl) or PEA116 (points = mean per well) (t = 2.950, p = 0.0256, unpaired t-test, two-tailed, Vcl: n = 3 wells, mean of 60 cells analyzed, PEA116: n = 5 wells, mean of 100 cells analyzed). **D.** Proportion of cells that do not divide (points = mean per well) (p = 0.0179, non-parametric Mann-Whitney test, two-tailed, Vcl: n = 3 wells, mean of 60 cells analyzed, PEA116: n = 5 wells, mean of 100 cells analyzed). **E**. Number of clones per cell (points = mean per wells) (t = 1.478, p = 0.1899, unpaired t-test, two-tailed, Vcl: n = 3 wells, mean of 60 cells analyzed, PEA116: n = 5 wells, mean of 100 cells analyzed). **F**. Mean of cell cycle length (in hours) (points = mean per wells) (t = 0.8275, p = 0.4396, unpaired t-test, two-tailed, Vcl: n = 3 wells, mean of 60 cells analyzed, PEA116: n = 5 wells, mean of 100 cells analyzed). **G**. Proportion of aNSPC that incorporated BrdU (t = 7.652, p = 0.0003, unpaired t-test, two-tailed, Vcl: n = 3 wells, mean of 60 cells analyzed, PEA116: n = 5 wells, mean of 100 cells analyzed). **H**. Confocal micrograph of cells immunostained for Caspase-3. **I.** Proportion of Caspase-3 (Casp3^+^) positive cells over the total population of aNSPC (DAPI^+^) after 24h exposure to Vcl, PEA116 or staurosporine (sts) (F = 2098, p < 0.001, One-Way ANOVA followed by Tukey’s multiple comparisons test, Vcl: n = 7 wells, PEA116: n = 12 wells, sts: n = 4 wells). **J**. Confocal micrograph of a hippocampal slice immunostained for Caspase-3 (red). **K.** Number of Casp3^+^ cells in the DG of mice after 8 days of treatment (t = 0.8415, p = 0.4197, unpaired t-test, two-tailed, n = 6 mice per group). **L.** Experimental design of the study of the effect of the PEA116 peptide on aNSPC quiescence in vivo. **M**. Confocal micrograph of a hippocampal slice immunostained for CldU (green) and IdU (red). **N-P**. Number of cells that are CldU^+^IdU^-^ (t = 0.5984, p = 0.5629, unpaired t-test, two-tailed, n = 6 mice per group) (**N**), CldU^+^IdU^+^ (t = 0.4932, p = 0.6325, unpaired t-test, two-tailed, n = 6 mice per group) (**O**) and CldU^-^IdU^+^ (t = 2.447, p = 0.0344, unpaired t-test, two-tailed, n = 6 mice per group) (**P**) in the DG of mice. **Q.** Percentage of PEA116-HA^+^ aNSPC over time after exposure. **R**. Percentage of aNSPC that internalized PEA116-HA after 15 minutes (F = 12.77, p =<0.0341, One-Way ANOVA followed by Tukey’s multiple comparisons test, n = 2 replicates). **S**. Fluorescence microscopy images of aNSPC with cytoplasmic internalization of PEA116-HA (red). **T.** Percentage of aNSPC that incorporated PEA116-HA in the cytoplasm (F = 335.1, p = 0.0003, One-Way ANOVA followed by Tukey’s multiple comparisons test, n = 2 replicates). **U.** Percentage of cells with nuclear or cytoplasmic localization of PEA116-HA. **V**. Schematic illustration of the workflow analysis of the two immunoprecipitation experiments. Results of the gene enrichment of the 264 interesting proteins found in the IPs. The three major processes that involved the greatest number of genes are represented. **W.** Proportion of pERK over ERK fluorescence intensity in HEK cells (F = 74.46, p < 0.001, One-Way ANOVA followed by Dunnett’s multiple comparisons test, Vcl: n = 2 replicates, PEA116: n = 5 wells (from 2 replicates), U0126: n = 2 replicates). **X, Y.** mRNA expression of NOTCH1 (t = 2.877, p = 0.0096, unpaired t-test, two-tailed, Vcl: n = 8 wells, PEA116: n = 13 wells) (**Y**) and HUWE1 (t = 3.215, p = 0.0046, unpaired t-test, two-tailed, Vcl: n = 8 wells, PEA116: n = 13 wells) (**Z**) in aNSPC. Histograms show average ± SEM. ns.: not significant p>0.05; *p<0.05; **p<0.01; ***p<0.001.

To assess the effect of PEA116 on apoptosis *in vitro*, cells were immunostained for Caspase-3 after treatment with PEA116, vehicle or staurosporine, which is known to induce apoptosis. While staurosporine substantially induced apoptosis, no difference in the proportion of Caspase-3^+^ cells were observed between vehicle– and PEA116-treated cells (**Fig. 3H, I**. Vehicle: 0.048 ± 0.002; PEA116: 0.044 ± 0.002, STS: 0.53 ± 0.02 Casp3^+^/total cells (%)). Similarly, *in vivo*, 8 days injections of PEA116 did not modify Caspase-3 staining (**Fig. 3J, K**. Tyrode: 0.75 ± 0.17; PEA116: 0.99 ± 0.2 Casp3^+^ cells/slice). Thus, although a reduction of cell death by PEA116 was observed in live-cell imaging, this effect could not be confirmed using Caspase-3 immunostaining *in vitro* and *in vivo*, suggesting that the reduction of cell death was likely not sufficient to explain the effect of PEA116 on adult neurogenesis.

Next, we assessed the effect of PEA116 treatment on cell quiescence *in vivo*. To this aim, we performed a dual thymidine analog injection using CldU and IdU^24,25^ in which mice received first three injections of CldU over a time window at 9 hours intervals. One hour after the last CldU injection, they received 3 injections of either Tyrode’s solution or PEA116 with 24 hours intervals, followed by three injections of IdU with 9 hours intervals, to label a secondary cohort of dividing cells. One hour after the last IdU injection, mice were sacrificed and perfused (**Fig. 3L**). Immunostaining for CldU and IdU revealed cells that expressed CldU only (cells that entered quiescence after PEA116 exposure), IdU only (cells that exited quiescence) and both CldU and IdU (highly proliferative cells, **Fig. 3M**). PEA116 did not increase the number of CldU^+^IdU^-^ cells (**Fig. 3N**. Tyrode: 3784 ± 195; PEA116: 3541 ± 356.8 cells/DG) or CldU^+^IdU^+^ cells (**Fig. 3O**. Tyrode: 1701.2 ± 110.1; PEA116: 1622.331 ± 115.1 cells/DG) but increased the number of CldU^-^IdU^+^ cells (**Fig. 3P**. Tyrode: 1596 ± 60.42; PEA116: 2070 ± 184.1 cells/DG), indicating that PEA116 activated quiescent cells.

### PEA116 increases ERK2 phosphorylation and decreases the expression of quiescence genes

Next, we assessed the cellular localization of PEA116. To this aim, PEA116 was tagged with a Human influenza hemagglutinin (HA) and administered to aNSPCs *in vitro*. Cells were then fixed at different time points, immunostained against HA and examined using imaging flow cytometry. The proportion of PEA116-HA^+^ cells increased from 0 to 15 minutes, before declining after 1h and being almost undetectable after 2h (**Fig. 3Q and Supp. Fig. 2J**. 0’: 0.3%, 10’: 2.6%, 15’: 31.5%, 1h: 10.9%, 2h: 0.14% of PEA116-HA^+^ cells). Furthermore, the incorporation of PEA-116-HA in aNSPC was cold-inhibited, suggesting that it resulted from an active process (**Fig. 3R**. 0’: 6.3 ± 0, 15’: 80 ± 20, 15’ ice: 22.33 ± 0.55 % of PEA116-HA^+^ cells). Finally, most cells incorporated PEA116 in the cytoplasm (**Fig. 3 S, T**. 0’: 0.052%, 15’: 73.94% ± 0.09, 1h: 71.64 ± 3.973 % of cytoplasmic PEA116-HA^+^ cells) and virtually none in the nucleus (**Fig. 3U**. Cytoplasm: 0’: 0%, 15’: 53.61%, 1h: 63.34%. Nucleus: 0’: 0%, 15’: 3.98%, 1h: 1.18% of internalized PEA116-HA^+^ cells). Thus, PEA116 was incorporated into the cytoplasm of aNSPC within minutes after exposure but did not cross the nuclear membrane.

To identify the molecular interactors of PEA116, we used immunoprecipitation. To this aim, we treated aNSPC with either vehicle or PEA116-HA (50 nM) for 30 minutes, lysed the cells and analyzed the immunoprecipitated proteins using LC/MS. We identified a total of 2339 immunoprecipitated proteins. From those, 472 proteins were either at least 4-fold more abundant in PEA116 than in vehicle condition, or completely absent from vehicle and possessed a spectral count greater than 5 in the PEA116 condition. A gene ontology analysis using Uniprot database identified 264 proteins that were restricted to the cytoplasm/cytosol (**Fig. 3V**). These included proteins involved in cell division, cell cycle, regulation of translation (**Supp. Fig. 3K**). Fourteen proteins were also identified by the Mammalian Adult Neurogenesis Gene Ontology (MANGO) database^26^ to play a role in adult neurogenesis. Two of these proteins are involved in stem cell quiescence: HUWE1, an E3 ubiquitin ligase that regulates protein stability within the Notch pathway^27^, and the Integrin-linked kinase (ILK) which, when expressed, promote cell quiescence^28^. Importantly, PEA116 immunoprecipitated with MAPK1 (ERK2), which plays a role in numerous cellular processes including stem cell proliferation and differentiation^29^. Of note, PEA116 is part of the c-terminal ERK-binding domain of PEA15, which regulates ERK phosphorylation and nuclear translocation^30^. To assess the effect of PEA116 on ERK2 phosphorylation, the ratio of pERK1/2/ERK1/2 was examined 10 to 90 minutes after PEA116 treatment, using an ERK phosphorylation ELISA assay kit. We found that PEA116 increased ERK1/2 phosphorylation (**Fig. 3W**. Vcl: 1 ± 0; PEA116: 1.174 ± 0.038, U0126: 0.451 ± 0.037 pERK/ERK).

To explore PEA116-induced changes in gene expression, we designed a quantitative gene expression array (“QuantiGene Sample Processing Kit” (QP1013)) to assess pathways involved in aNSC proliferation and quiescence. Four hours after aNSPC exposure to PEA116, cells were lysed and processed for hybridization to streptavidin-conjugated probes. We did not observe any difference in the expression of FMRP, GSK3B as compared to cells exposed to vehicle, neither did we find a difference in the expression of Hes1, Hes5, Ascl1, Fgfr3, Id4, Id3, ApoE or Aldoc (Data not shown). However, we observed a decrease in the mRNA expression of Notch1, which regulates the activity of the proactivation transcription factor Ascl1 (**Fig. 3X**. PEA116: 87.61 ± 2.6 % expression relative to control) and Huwe1 (**Fig. 3Y**. PEA116: 88.92 ± 2.3 % expression relative to control), consistent with a decrease in quiescence. Together, these results indicate that PEA116 enters the cytoplasm of aNSPC and interacts with several proteins involved in adult neurogenesis and decreases the expression of quiescence-inducing genes.

### PEA116 increases resilience to chronic stress

Finally, we assessed the functional implication of the increase in neurogenesis conferred by PEA116 treatment. An increase in adult neurogenesis has been shown to buffer stress response and reduce depression-like behaviors^7,8^. To test the functional impact of PEA116 on stress resilience, we first examined the effect of PEA116 injections on acute stress. Mice received injections of either Tyrode’s solution or PEA116 (5 mg/kg) for 8 consecutive days, followed by three injections of BrdU (100 mg/kg) the next day. Twenty-one days after the last injection, to enable new neurons to mature and functionally integrate into the hippocampal circuit^31,32^, an open field test (OFT) was conducted to evaluate basal anxiety levels. Mice were then subjected to an acute 6-hour restraint stress (ARS), followed by a second OFT, to assess stress-induced state anxiety (**Fig. 4A**). Anxiety scores were calculated based on time spent close to the walls and the center of the OFT, with values normalized and averaged to obtain scores between 0 and 1, where 1 indicates high anxiety and 0.5 represents standard non-anxious behavior. Before ARS, both groups showed normal values in anxious behavior (**Fig. 4B**. Anxious score Tyrode: 0.58 ± 0.05; PEA116: 0.57 ± 0.08), indicating that PEA116 did not display any effect on anxiety. Post-ARS, both groups exhibited increased anxiety levels, but no difference was observed between groups (**Fig. 4C**. Tyrode: 0.77 ± 0.04; PEA116: 0.72 ± 0.05 anxiety score). An assessment of adult neurogenesis confirmed that PEA116 administration increased adult neurogenesis as compared to vehicle (**Fig. 4D**. Tyrode: 339.7 ± 24.19; PEA116: 469.9 ± 59.66 BrdU^+^NeuN^+^ cells/DG). Thus, PEA116 did not confer resilience to ARS.

**Figure 4:**
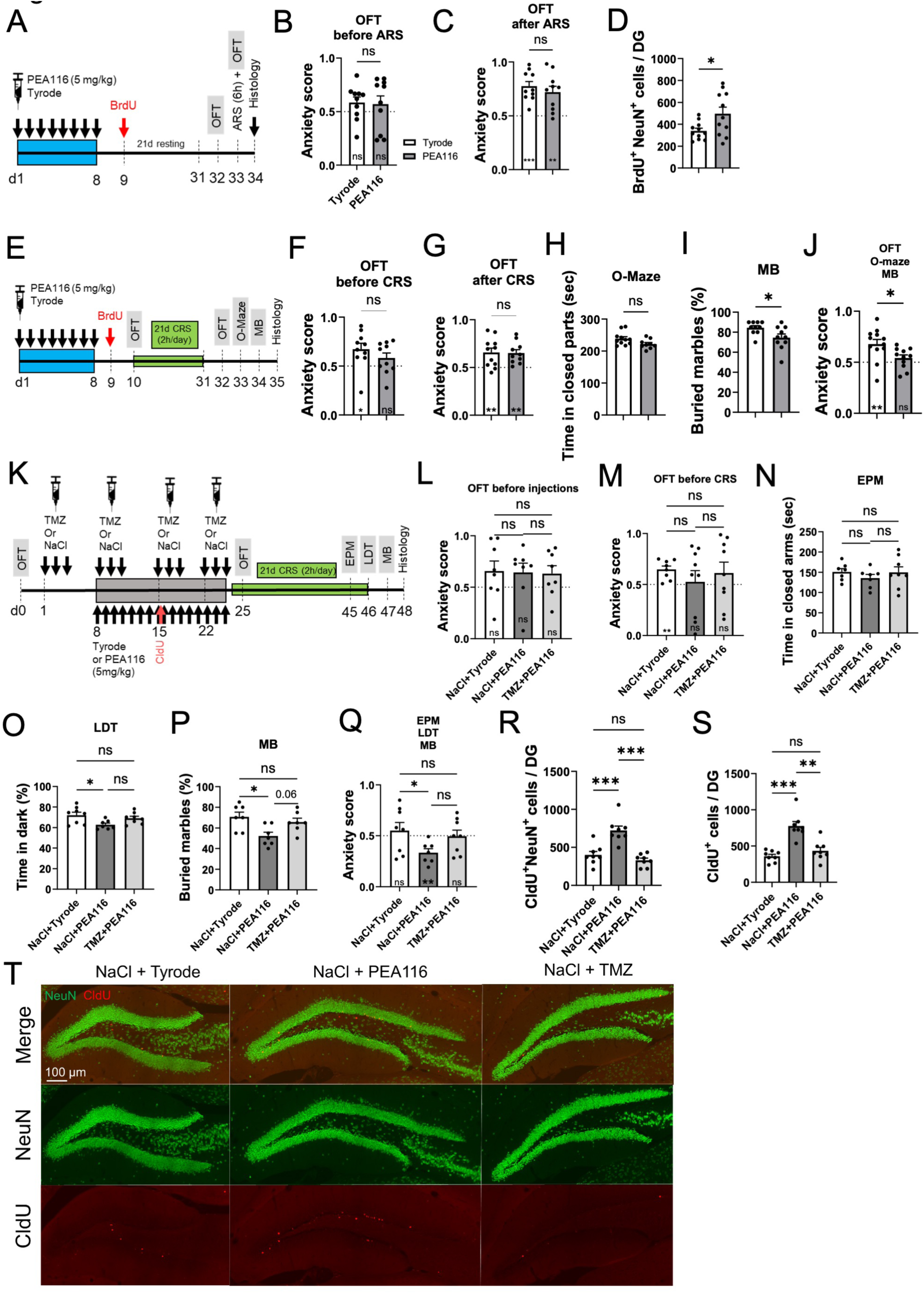
PEA116 increases stress resilience in a neurogenesis-dependent manner. **A.** Experimental design of the study of stress resilience to an Acute Restraint Stress (ARS) of mice treated with PEA116 or Tyrode (vehicle). **B-C.** Score of the time spent in the thigmotaxis and in the center of an OFT before ARS (**B**, t = 0.1634, p = 0.8720, unpaired t-test, two-tailed, n = 10 mice per group; Tyrode, PEA116 vs 0.5: t = 1.670, p = 0.1293; t = 0.9187, p = 0.3822, One-sample t-test) and after ARS (**C,** t = 0.7887, p = 0.4406, unpaired t-test, two-tailed, n = 10 mice per group; Tyrode, PEA116 vs 0.5: t = 6.364, p = 0.0001; t = 3.974, p = 0.0032, One-sample t-test). **D.** Number of newborn neurons in the DG (t = 2.443, p = 0.0240, unpaired t-test, two-tailed, n = 11 mice per group). **E.** Experimental design of the study of stress resilience to a Chronic Restraint Stress (CRS). **F-G.** Score of the time spent in thigmotaxis and in the center of an Open Field Test before the CRS (**F**, t = 1.141, p = 0.2687, unpaired t-test, two-tailed, Tyrode: n = 11 mice, PEA116: n = 10 mice; Tyrode, PEA116 vs 0.5: t = 2.880, p = 0.0182, t = 1.625, p = 0.1387, One-sample t-test) and after CRS (**G,** t = 0.1056, p = 0.9170, unpaired t-test, two-tailed, Tyrode: n = 11 mice, PEA116: n = 10 mice; Tyrode, PEA116 vs 0.5: t = 3.445, p = 0.0063, t = 4.250, p = 0.0021, One-sample t-test). **H.** Time spent in closed arms of an O-Maze (t = 2.052, p = 0.055, unpaired t-test, two-tailed, Tyrode: n = 11 mice, PEA116: n = 9 mice). **I.** Percentage of buried marbles in the Marble Burying test (MB) (t = 2.375, p = 0.0288, unpaired t-test, two-tailed, n = 10 mice per group). **J.** Combined anxiety score calculated from the OFT, OM and OMB tests (t = 2.441, p = 0.0236, unpaired t-test, two-tailed, Tyrode: n = 12 mice, PEA116: n = 11 mice; Tyrode vs 0.5: t = 3.895, p = 0.0025, PEA116 vs 0.5: t = 1.342, p = 0.2091, One-sample t-test). **K.** Experimental design of the study of the effect of TMZ on PEA116-induced stress resilience. **L-M.** Score of the time spent in the thigmotaxis and in the center of an Open Field Test (OLT) before injections (**L**, F = 0.02178, p = 0.9785, One-Way ANOVA followed by Tukey’s multiple comparisons test, n = 8 mice per group; NaCl+Tyrode, NaCl+PEA116, TMZ+PEA116 vs 0.5: t = 1.621, p = 0.1491, t = 1.587, p = 0.1566, t = 1.695, p = 0.1338, One-sample t-test) and before CRS (**M**, F = 0.4455, p = 0.6459, One-Way ANOVA followed by Tukey’s multiple comparisons test, NaCl+Tyrode: n = 7 mice, NaCl+PEA116: n = 8 mice, TMZ+PEA116: n = 8 mice; NaCl+Tyrode, NaCl+PEA116, TMZ+PEA116 vs 0.5: t = 4.424, p = 0.0031, t = 0.2313, p = 0.8229, t = 1.059, p = 0.3204, One-sample t-test). **N.** Time spent in closed arms of an EPM (F = 0.5779, p = 0.5706, One-Way ANOVA followed by Tukey’s multiple comparisons test, NaCl+Tyrode: n = 7 mice, NaCl+PEA116: n = 7 mice, TMZ+PEA116: n = 8 mice). **O.** Time spent in the dark zone of a LDT (F = 4.189, p = 0.0302, One-Way ANOVA followed by Tukey’s multiple comparisons test, NaCl+Tyrode: n = 8 mice, NaCl+PEA116: n = 7 mice, TMZ+PEA116: n = 8 mice). **P.** Percentage of buried marbles in the MBT (F = 5.716, p = 0.012, One-Way ANOVA followed by Tukey’s multiple comparisons test, n = n = n = 7 mice per group). **Q.** Anxiety score calculated from the combination of the EPM, LDT and MB tests (F = 3.038, p = 0.0704, One-Way ANOVA followed by Tukey’s multiple comparisons test, NaCl+Tyrode: n = 8 mice, NaCl+PEA116: n = 7 mice, TMZ+PEA116: n = 8 mice; NaCl+Tyrode, NaCl+PEA116, TMZ+PEA116 vs 0.5: t = 0.6547, p = 0.5336, t = 4.239, p = 0.0054, t = 0.07778, p = 0.9402, One-sample t-test). **R.** Number of CldU^+^NeuN^+^ newborn **S.** Number of CldU+ cells (F = 21.29, p < 0.001, One-Way ANOVA followed by Tukey’s multiple comparisons test, n = 8 mice per group) (F = 15.56, p = 0.0004, non-parametric Kruskal-Wallis test followed by Dunn’s multiple comparisons test, n = 8 mice per group)**. T.** Confocal micrographs of hippocampal slices showing newborn neurons (CldU^+^NeuN^+^). Histograms show average ± SEM. ns.: not significant p>0.05; *p<0.05; **p<0.01; ***p<0.001. To assess anxiety withing each group, comparisons between group mean and the hypothetical value of 0.5 are shown within each histogram bar. One-sample t-test. ns: non-significative, *: p<0.05; ** p<0.01.

Next, we tested the effect of PEA116 treatment on chronic restraint stress (CRS), consisting of restriction in a 50 ml Falcon tube for 2 hours each day for 21 consecutive days. Mice received injections of either Tyrode’s solution or PEA116 (5 mg/kg) for eight consecutive days, followed by three injections of BrdU (100 mg/kg) the following day. An OFT was conducted to evaluate basal anxiety levels, followed by 21 days of CRS and anxiety assessment using the OFT, O-Maze and Marble Burying test (MB, **Fig. 4E**). Before the CRS, there was no inter-group difference in anxiety (**Fig. 4F**. Tyrode: 0.47 ± 0.03; PEA116: 0.43 ± 0.03 anxiety score). After stress, both groups exhibited increased anxiety-levels in the OFT, but with no inter-group difference (**Fig. 4G**. Tyrode: 0.66 ± 0.04; PEA116: 0.65 ± 0.04 anxiety score). However, in the O-Maze, mice injected with PEA116 tended to spend less time in the closed sections, albeit not significantly (**Fig. 4H**. Tyrode: 0.55 ± 0.06; PEA116: 0.39 ± 0.05 seconds in closed segments)) and they buried significantly less marbles than control mice in the MB test (**Fig. 4I**. Tyrode: 84.50 ± 2.03; PEA116: 74.50 ± 3.69 % of buried marbles). When values from the OFT, O-Maze, and MB tests were normalized and combined to calculate a global anxiety score, mice injected with PEA116 displayed reduced anxiety following chronic stress as compared to control mice (**Fig. 4J**. Tyrode: 0.678 ± 0.05; PEA116: 0.541 ± 0.03 anxiety score). Thus, PEA116 increased resilience to CRS.

To assess whether the increase in adult neurogenesis was essential for the effects of PEA116 on resilience to CRS, we used Temozolomide (TMZ), a cytostatic drug that inhibits adult neurogenesis^33,34^. Mice were injected either with vehicle, PEA116 or PEA116 + TMZ. To optimize the effect of PEA116, mice were injected for 2 weeks and CldU was injected after one week of PEA116 injections. All mice underwent an OFT to assess baseline anxiety levels before and after injections. Mice were then subjected to CRS and anxiety was assessed using the elevated plus maze (EPM), light-and-dark test (LDT), and MB test (**Fig. 4K**). There was no inter-group difference in trait anxiety before and after injections (Anxiety score before injections, **Fig. 4L**. NaCl+Tyrode: 0.65 ± 0.1; NaCl+PEA116: 0.64 ± 0.09, TMZ+PEA116: 0.63 ± 0.08; After injections: **Fig. 4M**. NaCl+Tyrode: 0.64 ± 0.03; NaCl+ PEA116: 0.52 ± 0.1, TMZ+PEA116: 0.61 ± 0.11). After CRS, there was no inter-group difference in the time spent in the closed arm of the EPM (**Fig. 4N**. NaCl+Tyrode: 150.9 ± 8.2; NaCl+ PEA116: 135.7 ± 8.9, TMZ+PEA116: 149.7 ± 13.8 seconds). However, mice injected with PEA116 spent less time in the dark chamber in the LDT as compared to the vehicle group, an effect that was reduced in the PEA116-TMZ group (**Fig. 4O**. NaCl+Tyrode: 72.03 ± 2.91; NaCl+ PEA116: 62.69 ± 1.64, TMZ+PEA116: 69.04 ± 1.951 % time in the dark compartment). Similarly, in the MB test, mice injected with PEA116 buried fewer marbles than the control and PEA116-TMZ groups (**Fig. 4P**. NaCl+Tyrode: 70.7 ± 4.5; NaCl+ PEA116: 52.1 ± 3.76, TMZ+PEA116: 65.71 ± 3.69 % of buried marbles). When the values from the EPM, LDT, and MB tests were normalized and combined to calculate a global anxiety score, mice injected with PEA116 displayed reduced anxiety following chronic stress compared to control mice and this effect was reduced in the TMZ+PEA116 group (**Fig. 4Q**. Anxiety score: NaCl+Tyrode: 0.55 ± 0.08; NaCl+ PEA116: 0.33 ± 0.04, TMZ+PEA116: 0.49 ± 0.06 anxiety score). After the behavioral assessment, we assessed adult neurogenesis (**Fig. 4R-T**.). We found an increase in the number of CldU^+^ and CldU^+^-NeuN^+^ cells in mice injected with PEA116 as compared to the 2 other groups (**Fig. 4R**. CldU^+^cells/DG: NaCl+Tyrode: 359 ± 27.33; NaCl+PEA116: 777.1 ± 60.9, TMZ+PEA116: 434.2 ± 47.46; **Fig. 4S**. CldU^+^-NeuN^+^cells/DG: NaCl+Tyrode: 400.5 ± 45.55; NaCl+ PEA116: 722.7 ± 57.40, TMZ+PEA116: 329.6 ± 28.61).

Thus, PEA116 treatment increased adult neurogenesis and resilience to chronic-restraint stress and the effect on stress resilience was inhibited upon inhibition of adult neurogenesis.

## Discussion

In this study, we identified PEA116 a peptide produced by astrocytes that stimulates aNSC proliferation. PEA116 was found in the mouse and human brain and is derived from the PEA15 protein. *In vitro*, PEA116 quickly entered the cytoplasm of aNSPC where it interacted with several proteins involved in adult neurogenesis and induced ERK2 phosphorylation. Furthermore, PEA116 reduced the expression of the quiescence genes NOTCH1 and HUWE1, resulting in decreased aNSC quiescence. PEA116 administration resulted in increased number of Tbr2 and DCX-expressing progenitors and increased net neurogenesis. Finally, PEA116 administration increased resilience to chronic restraint stress, an effect that was dependent on its effect on adult neurogenesis.

Astrocytes play an essential role in the neurogenic niche and are known to regulate multiple steps of adult neurogenesis, including cell proliferation, differentiation, neuronal maturation, neuronal survival and synaptic integration^35,36,9,37^. Astrocytes of the DG in particular, have been found to be a very heterogenous population, which likely underlie the variety of functions in which they play a role^38^. We recently found that lactate secreted by astrocytes has antidepressant properties in a corticosterone model, an effect that is mediated by adult neurogenesis^39,40^. Similarly, adenosine 5’-triphosphate (ATP) released by astrocytes stimulates precursor proliferation via P2Y1-PLC-phosphatidylinositol 3-kinase (PI3K) signaling^41^ and has antidepressant properties^42^. Astrocytes also produce Wnt ligands that bind Frizzled/LRP heteromeric receptors expressed in adult hippocampal progenitor cells and regulate adult neurogenesis^43^ and have an impact on hippocampal-dependent memory performances^44^. FGF2 is primarily secreted by astrocytes and acts as a proliferative-inducing factor for precursors^45,46,47^. Interestingly, a decrease in astrocyte-derived FGF2 expression has been observed in aged mice, correlating with reduced neurogenesis in the aging hippocampus^48^. Moreover, studies have revealed astrocyte-secreted molecules such as thrombospondin-1 (TSP1), neurogenesin-1, IL-1β, IL-6, insulin-like growth factor binding protein 6 (IGFBP6), enkephalin, and decorin modulate neuronal fate and progenitor cell differentiation^36^. Although it remains unclear how these different signaling mechanisms are coordinated to maintain homeostasis in adult neurogenesis, these studies highlight the role of astrocytes as major regulators of adult neurogenesis. Furthermore, since astrocytic vesicular release is regulated by neuronal activity, astrocytes enable an on-demand production of new neurons in the DG.

Interestingly, astrocytes express glucocorticoid receptors, and chronic exposure to glucocorticoids changes the expression of genes involved in cell-cell signaling^49^. Thus, impaired astrocytic function after prolonged stress may contribute to the reduced adult neurogenesis observed in these conditions, an effect that may be alleviated by the administration of molecules that they would normally release, such as PEA116.

The PEA116 peptide is derived from the C-terminal portion of the PEA-15 protein (SEEEIIKLAPPP, aa 116-127). The PEA-15 protein encompasses two microtubule-binding regions (aa 98-107 and 122-129), which play a role in the stability of tubulins and may impact astrocyte morphology^50^. In addition, the PEA-15 protein sequesters the ERK1/2 kinase in the cytoplasm^51^ which, when released, can translocate to the nucleus and phosphorylate numerous targets to promote cell proliferation and differentiation^52^. Interestingly, the structural analysis of ERK2 bound to PEA-15 revealed that PEA-15 targets the 2 main ERK docking sites through a regulatory DEF docking site, at the N-terminal end of PEA-15 (aa 1-30 and 37-86) and a low affinity D-peptide docking site located at the C-terminal end of PEA-15 (aa 122-127) which is contained in the PEA116 peptide. Phosphorylation at Serine 104 and 116 enable the release of PEA-15 from ERK that becomes immediately activated^30^. Although it remains unclear whether PEA116 interferes with the interaction between endogenous PEA15 and ERK2, we found here that PEA116 interacts with ERK2 (**Supp. Fig.2K**) and increases its phosphorylation (**Fig. 3W**). This PEA116-ERK2 interaction may mediate cell cycle entry of quiescent aNSC, resulting in increased adult neurogenesis. However, the immunoprecipitation assay revealed that PEA116 also interacts with several other molecules involved in the regulation of adult neurogenesis, including cell division and quiescence. Thus, it remains unclear whether the effect of PEA116 on aNSC quiescence is mediated by its interaction with ERK2 or involves several pathways.

Together, these results identify PEA1126 as a peptide secreted by astrocytes that increases adult neurogenesis and resilience to chronic stress. This study highlights the contribution of astrocytes in the regulation of aNSC quiescence, a mechanism that is relevant to mood disorders.

## Material and methods

### Animals

Experimental protocols were approved by the Cantonal Veterinary Authorities (Vaud, Switzerland) and carried out in accordance with the European Communities Council Directive of 24 November 1986 (86/609EEC). All mice were adult male of 8 weeks-old at the beginning of the experiment, unless indicated. They were housed in a 12h light/dark cycle with free access to food and water and controlled temperature (23°C +-1°C) conditions. C57Bl/6J mice were purchased from Janvier (le Genest Saint Isle, France) or Charles River laboratories (Wilmington, MA, USA). The dnSNARE mice were a kind gift of P. Haydon (Tufts University, Boston, USA) and were generated by crossing the GFAP-tTA line expressing the Tet-Off tetracycline transactivator under the control of the GFAP promoter and the tetO-dnSNARE line containing a Tet-operator-regulated dominant-negative soluble N-ethylmaleimide-sensitive factor attachment protein receptor (dnSNARE) domain of synaptobrevin 2 as well as the reporter genes lacZ and GFP^18^. Doxycycline (dox) was administered in the food pellets (40 ppm, Harlan Laboratory Wi, USA) for dnSNARE and control animals (Wild Type (WT) and tetO-dnSNARE littermate) starting from mating of the parental pair onwards. Mice designated as runners were accommodated in standard cages containing a running wheel (Fast-Trac; Bio-Serv, USA) for a duration of 3 weeks, during which they had unrestricted access to the running wheel. Conversely, non-runner mice were housed in comparable cages positioned nearby but lacking a running wheel. Throughout all experiments, all mice were kept under a consistent 12-hour light/dark cycle at a controlled temperature of 22°C. Food and water were provided ad libitum.

### Cell culture

#### aNSPC

Mouse aNSPC were isolated from the dentate gyrus of adult Fisher 344 rats and cultured in medium DMEM/F12 + Glutamax (Gibco/Life Technologies 31331-028) supplemented with 1% of N2 Supplement 100x (Gibco/ Life Technologies 17502-048), Penicillin-Streptomycin 100x (PS, 50 UI/ml) (Gibco/Life Technologies 15140-122) and FGF-2 (20ng/ml) (PreProtech AF-100-18B) as previously described^53^. Mouse aNPCs were isolated from the dentate gyrus of adult C57Bl6/J mice and cultured in medium DMEM/F12 supplemented with 1% of N2 Supplement 100x, PS 100x (50 UI/ml), FGF-2 (20ng/ml) and EGF (20ng/ml) (PreProtech AF-100-15). ANSPCs were cultured on 10 cm dishes coated with Poly-Ornithine (10 mg/ml) (Sigma-Aldrich P3655), Poly-D-lysine hydrobromide at 0.1mg/ml (Sigma-Aldrich P6407) and Laminin at 20µg/ml (Gibco/Life Technologies 23017-015) for expansion and in 24-wells plates for experiments, with the same coating.

#### HEK-293

HEK-293 cell line was cultured in 10 cm dishes for expansion and in 24-wells plates for experiments in Dulbecco’s Modified Eagle Medium (DMEM) supplemented with 10% Horse serum and PSF 100x (50 UI/ml). Cells were grown in a humidified 5% CO2 incubator at 37°C.

#### Astrocytes

Astrocytes were prepared from postnatal day 1-3 mice (WT or dnSNARE mice) as previously described^17^. Briefly, mice were decapitated, and brains were rapidly collected. Meninges were removed, and the hippocampi and cortices were dissected. Tissues were mechanically triturated for homogenization and seeded onto 25 cm^2^ flasks in DMEM supplemented with Glutamax (Gibco/Life Technologies 10569-010), 10% of horse serum (Gibco/Life Technologies 16050-122) and PSF 100x (50 UI/ml) (Gibco/Life Technologies 15140-122). Cells were grown for one week in a humidified 5% CO2 incubator at 37°C. Every 2 days during the week, flasks were shaken to separate microglia from astrocytes and were washed with HBSS (Gibco/Life Technologies 14025-092). The enrichment of astrocytes in cultures was assessed by immunostaining. 98% of the cells expressed GFAP and none of the cells expressed the neuronal markers NeuN, DCX, TUJ1 or Map2 (data not shown), showing that this process efficiently removes neurons.

### Astrocyte conditioned medium (ACM) or astrocyte conditioned solution (ACS)

For *in vitro* experiments, ACM was prepared from confluent WT or dnSNARE astrocyte cultures in 25 cm^2^ flasks. Medium was removed and the cultures were washed twice with HBSS (Gibco/Life Technologies 14025-092) and replaced by aNSPC medium for 24h before being applied on aNSPC. For *in vivo* injections, ACS was prepared from confluent WT astrocyte cultures in 25 cm^2^ flasks. Medium was removed and the cultures were washed twice with HBSS (Gibco/Life Technologies 14025-092) to eliminate residual serum and replaced by 2 ml of Tyrode’s solution (HEPES 10 mM, NaCl 145 mM, KCl 5.4 mM, CaCl2 1.8 mM, MgCl2 0.8 mM, Glucose 10 mM, ddH2O qsp) for 24h. ACS was then collected and filtered with a 0.22 um filter (Millipore) to eliminate cellular debris. ACS was prepared fresh every day. For LC/MS analyses, astrocytes were grown in flasks for at least 10 days. Before the experiment, astrocyte cultures were washed twice with HBSS (Gibco/Life Technologies 14025-092) and the astrocyte medium was exchanged with 2 ml of Tyrode’s solution per flask and astrocytes cultures were incubated at 37°C for 24h. The medium was then collected and filtered with a 3 kDa filter (Amicon Ultra-4 centrifugal Filter 3 kDa MWCO, Merck UFC8003243). Five distinct preparations of ACS <3kDa were prepared from five astrocytes cultures and were then sent to the Protein Analysis Facility of the University of Lausanne for Mass Spectrometry analysis.

### Size exclusion filtration and Proteinase K treatment

ACS was prepared from astrocyte cultures with aNSPC medium and filtered with a 3 kDa filter (Amicon Ultra-4 centrifugal Filter 3 kDa MWCO, Merck UFC8003243 to obtain a < 3 kDa and a > 3 kDa fraction. To avoid depleting the medium from necessary molecules, these fractions were then supplemented with their opposite fraction of non-conditioned medium. ACS fractions were treated with Proteinase K (100 ug/ml) (Sigma-Aldrich P4850) during 3h at 37°C.

### Mass Spectrometry

#### Proteomics analyses

Five distinct preparations of ACS <3kDa were prepared from distinct astrocytes cultures. All samples were filtered using Amicon 0.5 ml Ultracel-10K filters (Millipore), and the eluates were acidified with TFA 20%, desalted on a StageTip C18 10µl (Thermo Scientific), speed-vacuum dried and redissolved in loading buffer. The prepared sample was injected into a Fusion Tribrid Orbitrap mass spectrometer using a 65-minute gradient with high-resolution accurate-mass MS/MS. The raw files were analyzed using Mascot and PEAKS software for peptide identification through both de novo and database search, with no enzyme specified in the search parameters

For immunoprecipitation experiments, samples were loaded on a 12 % mini polyacrylamide gel, migrated about 2.5 cm and stained by Coomassie. Gel lanes between 10-300 kDa were excised into 5 pieces and digested with sequencing-grade trypsin as described^54^. Extracted tryptic peptides were dried and resuspended in 0.05% trifluoroacetic acid, 2% (v/v) acetonitrile as loading buffer for Mass Spectrometry analyses.

#### Liquid Chromatography-Mass Spectrometry analyses

Data-dependent LC-MS/MS analyses of samples were carried out either on a Fusion Tribrid Orbitrap instrument (astrocyte, immunoprecipitation and microdialysate samples) or on a QExactive Plus mass spectrometer (immunoprecipitation samples), interfaced through a nano-electrospray ion source to an Ultimate 3000 RSLCnano HPLC system (Thermo Fisher Scientific). Peptides were loaded onto a trapping microcolumn Acclaim PepMap100 C18 (20 mm x 100 μm ID, 5 μm, Dionex) before separation on a reversed-phase custom packed 45 cm C18 column (75 μm ID, 100Å, Reprosil Pur 1.9 um particles, Dr. Maisch, Germany) with a 4-76 % acetonitrile gradient in 0.1% formic acid (total time: 65 min).

In Fusion instrument full MS survey scans were performed at 120’000 resolution. A data-dependent acquisition method controlled by Xcalibur software (Thermo Fisher Scientific) was used that optimized the number of precursors selected (“top speed”) of charge 2+ to 5+ while maintaining a fixed scan cycle time. Peptides were fragmented by higher energy collision dissociation (HCD) with a normalized energy of 32%. The precursor isolation window used was 1.6 Th, and the MS2 scans were done either at low resolution in the ion trap (immunoprecipitation and microdialysate samples) or in the orbitrap at 15’000 resolutions (astrocyte samples). The m/z of fragmented precursors was then dynamically excluded from selection during 60 s.

For microdialysate analyses, PEA116 peptide standard was also injected on the MS instrument, and additional targeted MS and MS/MS scans of m/z 661.865, corresponding to PEA116 peptide mass, were included in the LC-MS/MS method.

In QExactive instrument full MS survey scans were performed at 70,000 resolutions. In data-dependent acquisition controlled by Xcalibur software, the 10 most intense multiply charged precursor ions detected in the full MS survey scan were selected for higher energy collision-induced dissociation (HCD, normalized collision energy NCE=27 %) and analysis in the orbitrap at 17’500 resolution. The window for precursor isolation was of 1.5 m/z units around the precursor and selected fragments were excluded for 60s from further analysis.

#### Data processing

MS data of immunoprecipitation samples were analyzed using Mascot 2.7 (Matrix Science, London, UK) set up to search the rat (Rattus norvegicus) reference proteome based on the UniProt database (www.uniprot.org, version of November 5th, 2019, containing 29’958 sequences) and a custom contaminant database containing the most usual environmental contaminants and enzymes used for digestion (keratins, trypsin, etc.). Trypsin (cleavage at K,R) was used as the enzyme definition, allowing 2 missed cleavages. Carbamidomethylation of cysteine was specified in Mascot as a fixed modification. Protein N-terminal acetylation and methionine oxidation were specified as variable modifications. Mascot was searched with a parent ion tolerance of 10 ppm and a fragment ion mass tolerance of 0.02 (QExactive data) or 0.5 Da (Fusion data). Scaffold 4 (Proteome Software Inc., Portland, OR) was used to validate MS/MS based peptide and protein identifications.

MS data of astrocyte samples were analyzed using Mascot 2.6 ((Matrix Science, London, UK) set up to search the SwissProt database restricted to Mus musculus taxonomy (www.uniprot.org, version of November 2016, containing 16’846 mouse sequences), and a custom contaminant database containing the most usual environmental contaminants and enzymes used for digestion (keratins, trypsin, etc.). Nonspecific cleavage was used as the enzyme definition, and methionine oxidation was specified as variable modification. Mascot was searched with a parent ion tolerance of 10 ppm and a fragment ion mass tolerance of 0.02 Da. Scaffold 4 (Proteome Software Inc., Portland, OR) was used to validate MS/MS based peptide and protein identifications.

MS data of microdialysate samples were analyzed using Mascot 2.8 ((Matrix Science, London, UK) set up to search the SwissProt database restricted to Mus musculus taxonomy (www.uniprot.org, version of April 2021, containing 17’097 mouse sequences), and a custom contaminant database containing the most usual environmental contaminants and enzymes used for digestion (keratins, trypsin, etc.). Nonspecific cleavage was used as the enzyme definition. Protein N-terminal acetylation and methionine oxidation were specified as variable modifications. Mascot was searched with a parent ion tolerance of 10 ppm and a fragment ion mass tolerance of 0.5 Da. Scaffold 5 (Proteome Software Inc., Portland, OR) was used to validate MS/MS based peptide and protein identifications.

In Scaffold validation, peptide identifications were accepted if they could be established at greater than 90.0% probability by the Peptide Prophet^55^ or the Scaffold Local FDR algorithm. Protein identifications were accepted if they could be established at greater than 95.0% probability and contained at least 1 identified peptide. Protein probabilities were assigned by the Protein Prophet algorithm^55,56^. Proteins that contained similar peptides and could not be differentiated based on MS/MS analysis alone were grouped to satisfy the principles of parsimony. Proteins sharing significant peptide evidence were grouped into clusters.

### Peptide synthesis

Peptides P1, P2, P3, P4, PEA116 and PEA116-HA were synthesized by the Peptide & Tetramer Core facility (Epalinges (UNIL), Switzerland). Peptides were produced by Solid Phase Peptides Synthesis (SPPS) using fmoc/tBu strategy. Peptides were assembled using CSBio 136M instrument, purified by reverse-phase HPLC and analyzed by UHPLC-MS (Agilent Technologies). Identity was confirmed by Mass Spectrometry (+/-1 da of calculated theoretical peptide mass) and the purity of all peptides was above 95%, based on Abs at 215 nm. Peptides were kept lyophilized at –20°C or below and resuspended in water prior to use.

### scRNAseq analyses

Expression of PEA15 in single-cell RNAseq data from Hochgerner et al. 2018 and Franjic et al. 2021^19,20^ was analyzed using Seurat v4.1.1. Single-cell sequencing data and cell-type annotations of mouse and human DG cells were obtained from the GEO repository (GSE95315 and GSE186538). Basic filtering of cells based on the fraction of mitochondrial reads (< 6%) was performed using Seurat v4.1.1. Cells from the human dataset were furthermore filtered based on the number of available features (nFeature_RNA > 600 & nFeature_RNA < 12000). Normalization and scaling of the sequencing data was performed using the default settings of Seurat’s ‘NormalizeData’ and ‘ScaleData’ functions. Principal components were calculated by ‘RunPCA’. Dimensionality reduction was performed by ‘RunUMAP’, using the first thirty principal components as input.

### Analysis of human brain interstitial space proteomics

Human Brain extracts were obtained from the Lausanne Hospital Neurobanks (BB-0063), with prefrontal cortex tissue from healthy control (∼1g) cut into smaller pieces and digested in a Papain/Hibernate-E solution to release interstitial fluid while minimizing cell lysis as previously described^57^. After differential centrifugation (300g, 2,000g, and 10,000g), the supernatant was stored at –80°C. Extracellular vesicles were isolated following the MISEV 2018 guidelines^58^ using size exclusion chromatography (SEC), with an average of 7.94 x 10E10 vesicles/g of tissue recovered. F1-4 fractions were then subjected to a modified iST protein digestion method. Samples were mixed with miST lysis buffer (1% sodium deoxycholate, 100 mM Tris, 10 mM DTT), heated at 95°C, reduced with chloroacetamide, and digested with Trypsin/LysC for 1 hour, followed by a second digestion. Peptides were desalted using strong cation exchange, eluted in 80% acetonitrile, and analyzed using a high-resolution Orbitrap Exploris 480 mass spectrometer with C18 column separation and a 140-minute gradient. Mass spectrometry data were acquired with data-dependent acquisition and processed using MaxQuant, searching against the UniProt human reference proteome and contaminant database, with peptide and protein identifications filtered at 1% FDR as previously described^59^.

### *In vitro* experiments

#### Cell culture and immunostaining

Human astrocytes were purchased from Sciencell (Sciencell, 1800). ANSPC were plated at in a 24-well-plate coated at a density of 60 000 cells per well. For treatment with ACS, medium was removed and 500 µl of ACS was added on aNSPC for 24h. For treatment with the PEA116 or PEA116-HA peptide, the peptides were solubilized in ddH2O and aNSPC were treated with PEA116 at 50 µM in 500 µl of aNSPC medium or with PEA116-HA at 100 µM in 500 µl of aNSPC medium for 24h. DdH2O was used as control. To assess cell proliferation, the medium was supplemented with 5 μM BrdU for 30 min and washed twice with pre-heated fresh medium followed by 1h incubation. Cells were fixed with 4% paraformaldehyde for 30 minutes and rinsed with PBS 0.1M or fixed with ice-cold MeOH 100% for 15 minutes and rinsed with PBS 0.1M before p-ERK1/2 and ERK1/2 immunostaining. Immunostainings against p-ERK, ERK, RFP, and Caspase-3 were processed as follows: after 3 washes with PBS 0.1M, cells were saturated with blocking solution (PBS 0.1M, decomplemented Horse serum 10%, Triton 100x 0.3%) during 1h. Plates were incubated with primary antibodies rabbit anti-ERK1/2 (1/500, Cell Signaling 4695), rabbit anti-p-ERK1/2 (1:500, Cell Signaling 9101S), rabbit anti-RFP (1:500, Rockland 600-401-379), mouse anti-RFP (1:500, ThermoScientific MA5-15257), rabbit anti-HA (1:500, Cell Signaling C29F4) or rabbit anti-Caspase-3 (1:500, Cell Signaling 9579) overnight at 4°C. Fluorescence was achieved by adding secondary antibodies goat anti-mouse Alexa Fluor 488 (1:300, Invitrogen A11029), goat anti-rabbit AlexaFluor488 (1:300, Invitrogen A11034), goat anti-rabbit AlexaFluor594 (1:300, Life Technologies A11037) or goat anti-mouse AlexaFluor594 (1:300, Invitrogen A11032) during 1h at room temperature. Finally, cells were incubated with DAPI (1:1000) for 10 minutes to reveal their nuclei. Immunostaining against BrdU was processed as follows: after 3 washes with PBS 0.1M, aNSPC were treated with HCl 2N for 15 minutes at 37°C then rinsed with borate buffer 0.1M for 15 minutes. After 3 washes with PBS 0.1M, cells were saturated with blocking solution (PBS 0.1M, decomplemented Horse serum 10%, Triton 100x 0.3%) during 1h. Plates were incubated with primary antibodies mouse anti-BrdU (1:250, BdBiosciences B44/347580) overnight at 4°C. Fluorescence was achieved by adding secondary antibodies goat anti-mouse Alexa Fluor 488 (1:300, Invitrogen A11029) during 1h at room temperature. Finally, cells were incubated with DAPI (1:1000) for 10 minutes to reveal their nuclei.

#### Image acquisition and analysis

Images were acquired using an Eclipse Ti2 inverted microscope (Nikon). The number of BrdU-labeled aNSPC was counted in one randomly selected large field (tiles 4×4) in each well of the plate with Imaris software. At least 6 wells were analyzed by treatment. The number of BrdU-labeled aNSPC was compared with the total number of aNSPC in each selected field to obtain a proliferation ratio.

#### Live-cell imaging of aNSPC

ANSPC were plated in a 8-wells Ibidi plates (Ibidi 80826) at a density of 40’000 cells per well in SILAC Advanced DMEM/F-12 Flex Media (Gibco/Life Technologies A2494301) supplemented with 0.7mM L-arginine, 0.5mM L-lysine, 5mM glucose, N2 Supplement 100x (Gibco/ Life Technologies 17502-048), PSF 100x (50 UI/ml) (Gibco/Life Technologies 15140-122), FGF-2 (20ng/ml) (PreProtech AF-100-18B) and GlutaMAX™ Supplement 1X (Gibco/Life Technologies 35050061). Thirty minutes after plating, cells were placed in the live imaging incubation chamber of an Eclipse Ti2 inverted microscope (Nikon), at 37°C and 5% CO2. Recording started immediately, capturing basal proliferation for 4 hours in 4 spots per well, every 30 minutes. Five out of the eight wells were then treated with 50 µM PEA116 and recording continued for a total duration of 48 hours. Cells that remained in the imaging frame for the entire duration of the experiment, that did not form clusters, or lacked morphological stem cell features, were randomly selected for fate analysis, for a total of 60 control cells and 100 PEA116 cells.

#### Quantigene Plex Assay

On the first day, mouse aNSPC were plated in 24-wells plate at a density of 60’000 cells per well. One day later, cells were treated with 50 µM PEA116 for 4h, then processed for gene expression analysis according to the protocols of ThermoFisher (“QuantiGene Sample Processing Kit” (QP1013) and “QuantiGene™ Plex Gene Expression Assay, USER GUIDE”). Briefly, cells were lysed and hybridized with specific RNA target probes. Formed RNA-probe hybrids were captured onto magnetic beads or well plates, washed to remove nonspecific contaminants. Detection of the hybridized RNA was performed by measuring the streptavidin signal associated with the probes. RNA levels in the sample were quantified based on the detected signal and reference standards provided in the kit. A panel of 50 genes were chosen (AKT3, ALDOC, APC, APOE, ASCL1, AXIN1, BCL9, BTRC, CASP9, CCND1, CCNE, CDK4, CREB1, CTNNB1, CXCR4, DCX, DKK1, DVL1, E2F1, E2F3, FGFR3, FMR1, FZD1, GSK3B, HES1, HES5, HPRT1, HUWE1, ID3, ID4, LEF1, LMNB1, LRP5, MAPK1, MAPK2, MKI67, MTOR, MYC, NEUROD1, NEUROG1, NOTCH1, RB1, RPLP0, RPL27, RPS26, SFRP3, TCF7, TCF7L1, TP53, TUBB) and RNA of aNSPC were analyzed using Luminex MAGPIX™ instrument (ThermoFisher) and xPONENT™ software. A total of three biological replicates were made, with at least 4 replicates per treatment. The background values were subtracted from the data and were normalized by the values of averaged reference genes. The geometric mean was calculated for each treatment and control values for each gene and was normalized to 1 to allow comparison with PEA116-treated values.

#### FACS

The PEA116 peptide was first conjugated to an HA tag (YPYDVPDYASG-SEEEIIKLAPPP). On day 1, Rat aNSPC were plated in 6-wells plate coated with Poly-Ornithine (10 mg/ml) (Sigma-Aldrich P3655), Poly-D-lysine hydrobromide at 0.1mg/ml (Sigma-Aldrich P6407) and Laminin at 20µg/ml (Gibco/Life Technologies 23017-015) at a concentration of 200’000 cells per well. On day 3, cells were treated with the PEA116-HA [100 µM] for 0 minute (as control for no peptide), 15 minutes, 1h or 2h at 37°C. By the end of incubation with the PEA116-HA, cells were dissociated with TrypLE Express (Invitrogen 12604-013) and placed in 12×75mm Falcon tubes (Life Sciences 352052), washed with PBS 1X, centrifuged for 5 minutes at 300g at 4°C and supernatant was discarded. Cells were washed with Flow Cytometry Staining Buffer (500ml PBS 1X + 2.5g BSA (Sigma Aldrich A9418) + 3ml EDTA 0.5M (Gibco/Life Technologies 15575020)) by centrifugation at 300g for 5 minutes. Supernatant was discarded and cells were vortexed in residual solution (∼100µl). To analyze cytoplasmic compartment, the “Intracellular Fixation & Permeabilization Buffer Set” was used (eBioscience 88-8824). Cells were fixed with 100µl with IC Fixation Buffer (eBioscience 00-8222-49) for 30 minutes at room temperature and protected from light. Cells were washed after that 2ml of Permeabilization Buffer (eBioscience 00-8333-56) was added in the tubes at 600g for 5 minutes at room temperature. Cells were resuspended in a Permeabilization buffer containing conjugated primary antibody anti-HA AlexaFluor647 (1:500, Biolegend 16B12) and were incubated for 30 minutes at room temperature and protected from light. Cells were finally washed once with Permeabilization Buffer and once with Flow Cytometry Staining Buffer at 600g for 5 minutes and finally resuspend in Flow Cytometry Staining Buffer before analysis. To analyze nuclear compartment, we used the “Foxp3 / Transcription Factor Staining Buffer Set” (eBioscience 00-5523), the protocol was similar except that Fixation/Permeabilization solution (eBioscience 00-5223-56 and eBioscience 00-5123-43) was used to fix the cells and a DAPI was added to mark cell nuclei. The percentage of PEA116-HA positive cells were analyzed on a LSRII SORP (Becton Dickinson) machine with FACS Diva software version 8.0. The percentage of cytoplasmic and nuclear PEA116-HA positive cells were analyzed using an Amnis ImageStream X flow cytometer (Cytek) with IDEAS software version 6.3.

#### Immunoprecipitation

Rat aNPCs were plated at a confluency of 10^6^ cells/ml in a 6-wells coated plate in the morning of the experiment. In the afternoon, cells were treated with PEA116-HA [100µM] or vehicle for 30 minutes at 37°C. Cells were then washed twice with ice-cold HBSS (Gibco/Life Technologies 14025-092) and ice-cold RIPA (ThermoScientific 89900) supplemented with protease inhibitor cocktail (PIC, 1:100, Merck P8340) was added to lyse the cells before scraping and harvesting in lock tubes. Cells were incubated 15 minutes at 4°C on orbital shaker in RIPA buffer to finish lysis. Lysate was centrifuged at 13’000 rpm for 10 minutes at 4°C and the supernatant was transferred in a new tube. Cells were incubated with 2 µg rabbit anti-HA antibody (Cell Signaling C29F4) overnight at 4°C on orbital shaker. The next day, 40 µl of protein A/G linked to agarose beads (Santa Cruz SC-2003) was added in each tube and incubated overnight at 4°C under agitation. 24h later, the immunoprecipitate was centrifuged at 4°C and 2500 rpm for 5 minutes. The supernatant was transferred in a new tube and PBS containing 1/100 of PIC at 4°C was added. After 3 washes with PBS (500 µl) beads were resuspended in 40 µl of PBS. We used 15 µl of each solution to perform SDS-gel to visualize all the protein in each condition and for further Mass Spectrometry analysis (See “Mass Spectrometry”).

#### ERK phosphorylation

The level of phosphorylated ERK1/2 in HEK-293 cell cultures was determined using the “ERK Phosphorylation ELISA Assay Kit” (ABIN1019677). Cells were treated with PEA116 (100 µM) for 10, 20, 30, 60 and 90 minutes prior to fixation and analysis. The ERK1/2 inhibitor U0126 (20 µM, SigmaAldrich, 662005) was used as a negative control.

### In vivo experiments

#### Injections

Mice were injected intraperitoneally with 200 µl of ACS pre-warmed at 37°C, prepared from a different dish of astrocytes each day for 8 days. For the PEA116 peptide injection, mice were injected intraperitoneally with PEA116 dissolved in Tyrode’s solution at a concentration of 5 mg /kg each day for 8 days in all experiment except for the study of quiescence *in vivo,* where mice were injected 3 times over 48h. BrdU was dissolved in 0.9% NaCl and injected at a concentration of 100 mg/kg 3 times each 2h (Sigma-Aldrich, Buchs, Switzerland). CldU was dissolved in 0.9% NaCl and injected at a concentration of 42.5 mg/kg either 3 times each 9h for the experiment about quiescence exit *in vivo*, or 3 times each 2h to assess net neurogenesis. IdU was dissolved in a solution containing 20% NaOH 0.2M + 80% NaCl 0.9% at a concentration of 57.5 mg/kg and was injected 3 times every 9h. TMZ was dissolved in NaCl 0.9% and injected at a concentration of 25 mg/kg (Sigma-Aldrich, T2577).

#### Tissue collection and preparation

Mice received a lethal dose of pentobarbital (10 mL/kg, Sigma, Switzerland) via intraperitoneal injection. Following this, they were perfused with 0.9% NaCl for 3 minutes, followed by perfusion with 4% paraformaldehyde (PFA) for 3 minutes (Sigma-Aldrich, USA) dissolved in 0.1 M Phosphate Saline Buffer (PBS, pH 7.4). Their brains were dissected, fixed overnight at 4°C in PFA 4%, and then incubated for 24h in a 30% sucrose solution at 4°C (Sigma-Aldrich, USA). The brains were rapidly frozen using isopentane at –40°C. Forty μm thick coronal sections were prepared using a cryostat (Leica MC 3050S) and were preserved in cryoprotectant (30% ethylene glycol + 25% glycerin in 0.1 M PBS) at −20°C.

#### Immunochemistry and antibodies

Immunochemistry was performed on every 6th section of the dentate gyrus. Sections were washed 3 times in PBS 0.1 M and permeabilized/blocked using PBS 0.1 M containing 0.3% Triton-X 100 and 10% Horse serum. BrdU immunohistochemistry was preceded by DNA denaturation by incubation in formic acid 50% formamide/ 50% 2X SSC buffer (2X SSC is 0.3 M NaCl and 0.03 M sodium citrate, pH 7.0) at 60°C for 2 h, rinsed twice in 2X SSC buffer, incubated in HCl 2M for 30 min at 37°C and rinsed in 0.1 M borate buffer pH 8.5 for 15 minutes and 6 times in PBS 0.1M for 10 minutes each. Then, sections were incubated overnight at 4°C with one of the following primary antibodies: rat monoclonal mouse anti-BrdU/IdU (1:250 or 1:500, BdBiosciences B44/347580), rat monoclonal anti-BrdU/CldU (1:500, Abcam ab6326), rat anti-Tbr2 (1:250, ebioscience 14-4875-82), mouse anti-NeuN (1:500, Chemicon MAB377), rabbit anti-NeuN (1:1000, Abcam EPR12763), rabbit anti-GFAP (1:500, Dako Z0334), rabbit anti-Sox2 (1:500, Millipore AB5603), mouse anti-DCX (1:400, Santa Cruz sc8066), rabbit anti-DCX (1:500, Abcam ab18723), rabbit anti-Caspase-3 (1:500, Cell Signaling 9661S). Sections were then incubated for 1h15 at room temperature with the corresponding fluorescent secondary antibodies: goat anti-mouse AlexaFluor594 (1:300, Invitrogen A11032), goat anti-rat AlexaFluor488 (1:300, Invitrogen A11006), donkey anti-mouse AlexaFluor647 (1:300, Invitrogen A31571), donkey anti-rabbit AlexaFluor647 (1:300, Invitrogen A31573), goat anti-rat AlexaFluor594 (1:300, Invitrogen A11007), goat anti-rabbit AlexaFluor488 (1:300, Invitrogen A11034). Finally, sections were incubated in DAPI (1/1000) to reveal their nuclei and washed 3 times in PBS 0.1M before being mounted on slides.

#### Imaging

The number of BrdU-positive cells was counted in the subgranular region of the dentate gyrus. This was done using an epifluorescence microscope (Zeiss AxioSkop2Plus) equipped with a camera (Zeiss AxioCamMRm) and a 20x objective. All cells were quantified throughout the entire thickness of the section. Using ImageJ software, the contour of the dentate gyrus of each section was traced to calculate the volume of the DG and determine the ratio of BrdU-positive cells in the whole volume of the dentate gyrus. Sections marked with GFAP, Tbr2, Sox2, DCX, NeuN or Caspase-3 were imaged using a laser scanning confocal microscope (Zeiss LSM780 GaAsp) equipped with a 20x objective (numerical aperture 1.0) and an automated optical configuration optimized for sequential visualization of each fluorescence channel or using a Nikon NI-E spinning disk with 20x objective. For each tissue volume sampled, adjacent image planes through the z-axis were collected throughout the entire thickness of the tissue. Images were analyzed with ZenBlue software (Zeiss).

#### Histological analysis of inflammation

Immunochemistry was performed on every 6th section of the dentate gyrus. Sections were washed 3 times in PBS 0.1 M and blocking of non-specific binding was achieved by incubating in PBS 0.1 M containing 0.3% Triton-X 100 and 10% Horse serum. Then, sections were incubated over night at 4°C with the following primary antibodies: rat anti-CD68 (1:1000, Abcam 53444), rouse anti-GFAP (1:500, Millipore MAB360), rabbit anti-Iba1 (1:500, Wako 019-19741). Then, sections were incubated for 1h15 at room temperature with the following fluorescent secondary antibodies: goat anti-rat Alexa Fluor 488 (1:300, Life Technologies A11006), goat anti-rabbit Alexa Fluor594 (1:300, Life Technologies A11037), donkey anti-mouse Alexa Fluor 647 (1:300, Invitrogen A31571). Finally, sections were incubated in DAPI (1/1000) to reveal their nuclei and washed 3 times in PBS 0.1M before being mounted on slides. Images acquired with the Nikon Spinning Disk microscope (details provided) were analyzed using the NIS-Elements software. The General Analysis (GA) tool within NIS-Elements facilitated automated morphometric and density analysis. Masks were created based on Iba1 staining for microglia or GFAP coverage for astrocytes. Regions of Interest (ROIs) were delineated according to anatomical landmarks to extract information about the total dentate gyrus area. For detailed microglial morphometry, multipoint images were processed using IMARIS 5.1 (Oxford Instruments) for three-dimensional reconstructions. The filament tool in IMARIS enabled reconstruction of microglial fine processes, providing information on ramification.

#### Microdialysis

Mice were anaesthetized with isoflurane and were placed in a stereotaxic frame using a mouse adaptor (Kopf) with modified ear bars. Microdialysis probes were implanted in the DG (Bregma: –2 mm anteroposterior, 1.75 mm lateral and –2 mm dorsoventral). The active dialysis surface length of the cellulose membrane (concentric design, molecular weight cutoff 20 kDa) was 1 mm. The probe was secured in place with dental cement on the skull. Microdialysis experiments started 24 h after surgery. Ringer solution (125 mM NaCl, 2.5 mM KCl, 1.26 mM CaCl2, 1.18 mM MgCl2, 0.20 mM NaH2PO4) was perfused through the microdialysis probe at a flow of 1.2 µl/min using a high precision pump (CMA 400 syringe pump, CMA Sweden). Experiments were performed during the light period. The dialysates were collected 4 times every 15 minutes in small eppendorf tubes and directly placed at –20°C until the end of the procedure and then frozen at –80°. At the end of the experiment, animals were euthanized with an overdose of pentobarbital and brains were removed and stored in 4% paraformaldehyde until Mass Spectrometry analysis.

#### Chronic and acute restraint stress

The 21 days CRS was performed as previously described,^60^. Mice were placed headfirst into 50 ml Falcon tubes (11.5 cm in length, 3 cm in diameter) with perforations at the bottom for ventilation, and tissue was placed at the end to adjust the constraint to the size of the mouse, allowing the tail to extend out. This restraint lasted for 2h daily for 21 consecutive days. For the ARS, we used the same protocol as for the CRS, with a restraint of 6 hours. After this, mice were returned to their home cages for 20 minutes before undergoing an open-field test to evaluate anxiety-like behavior.

### Behavioral tests

#### Open-field test (OFT)

The OFT was conducted following previously established protocols. The setup comprised a square Plexiglas enclosure measuring 40 x 40 x 40 cm, illuminated with dimmed light (25 lux). Mice were placed facing the wall of the arena and given 10 minutes to freely explore the arena. The arena was divided into zones including a virtual center zone (15 x 15 cm) and a thigmotaxis zone (within 10 cm of each wall), which were analyzed to assess anxiety-like behavior. A video tracking system (Anymaze software) recorded the movement of each mouse, total distance travelled, and time spent in each zone. To prevent olfactory cues, the OF was cleaned after each trial.

#### Elevated plus maze test (EPM)

The apparatus was constructed from black PVC with a white floor and consisted of a central platform (5 x 5 cm) elevated 65 cm from the ground, featuring two open arms (30 x 5 cm) and two closed arms (30 x 5 x 14 cm) arranged in opposing pairs. Light intensity was maintained at 15 lux in the open arms and 4 lux in the closed arms. Animals were placed at the end of the closed arms facing the wall and allowed to freely explore the apparatus for 5 minutes. Anymaze software was used to follow the movements of the mice and to measure the time spent in each arm and at the risk zones (edges of the open arms). To prevent olfactory cues, the EPM was cleaned after each trial.

#### Light Dark Test (LDT)

A Plexiglas box measuring 60 x 40 x 21 cm was utilized, divided into a dark compartment (20 x 40 x 21 cm) and a light compartment (40 x 40 x 21 cm) illuminated at 400 lux. The compartments were separated by an open door (5 x 5 cm) positioned centrally at floor level. Each mouse was introduced into the light chamber facing the door and allowed to explore freely for 6 minutes. Anymaze software was used to follow the movements of the mice and to measure anxiety-like behavior by analyzing the time spent in each zone. To prevent olfactory cues, the chambers were cleaned after each trial.

#### Marble Burying test (MB)

The MB test was conducted following previously established protocols^61^. A cage measuring 40 x 25 x 20 cm was filled to a depth of approximately 5 cm with husk bedding material, which was evenly spread across the cage’s surface. Twenty plain dark glass marbles, each with a diameter of 1.4 cm, were then arranged in a 4 × 5 grid on top of the bedding. Each mouse was introduced into the cage and allowed to explore it for 20 minutes under 300 lux lighting conditions. Following the test, the mice were removed from the cage, and the number of marbles buried with bedding up to 2/3 of their depth was counted. Between each trial, the marbles were removed, and the bedding was stirred, spread, and compacted before the marbles were replaced.

### Anxiety scores

The anxiety score was computed following the methodology outlined in previous studies^24,25^, involving the averaging of standardized scores from various anxiety-related behavioral tests. Standardization was achieved by subtracting the minimum value of the entire population from each animal’s value and then dividing the result by the difference between the maximum and minimum values of the entire population: (x – min value) / (max value – min value). This process yielded scores between 0 and 1, with higher scores indicating greater levels of anxiety. The score was derived from factors such as the time spent in the dark chamber of the LDT and the time spent in the closed arm of the EPM. In OFT, it included the time spent in thigmotaxis and in the center of the arena.

### Statistics

All statistical tests were performed with GraphPad Prism (GraphPad 9 software, San Diego, CA, USA) or RStudio (RStudio Team (2023). RStudio: Integrated Development for R. RStudio, PBC, Boston, MA, Version 1.2.5033) using a critical probability of p < 0.05. Statistical analyses performed for each experiment are summarized in each figure legend with the chosen statistical test, sample size ‘n’ and p values. All values are given as mean ± S.E.M. For experiments with two groups, a Shapiro-Wilk test was performed before all tests to assess normality followed by an unpaired Student t-test or a non-parametric Mann-Whitney test according to the results of the normality test. In experiment with more than two groups, a Shapiro-Wilk test was performed to assess normality, and a Bartlett test of heteroscedasticity were done before all tests. One-Way ANOVA or Kruskal-Wallis test was performed according to the results of the pre-tests. A post-hoc Tukey’s or Dunn’s test was performed for multiple comparisons. In Figure 4, for OLT (Figure 4I, 4J), NOR (Figure 4K), OFT (Figure 4N, 4O, 4R, 4S) and the anxiety score combining EPM + LDT + MB (Figure 4W), the difference between groups was analyzed by an unpaired Student t-test and. The difference between each group and the theoretical mean for anxiety score set at 0.5 was assessed with a one sample t-test.

## Acknowledgements

The authors wish to thank Patrice Waridel for technical assistance with Mass Spectrometry analyses. Romain Bedel and Francisco Sala de Oyanguren for their help with FACS analyses. The authors also wish to thank Pr. Phil Haydon and his team for sharing the dnSNARE mouse and advice on breeding and doxycycline administration.

## Funding

This work was supported by the Swiss National Science Foundation (310030_201015) to NT. A.S. and R.B. were supported by grants from the German Research Foundation (DFG; BE5136/6-1 and Research Training group 2162 to R.B., Emmy Noether Program grant 455354162 to A.S.)

## Author contributions

NT, CC, FC and SS conceptualized the study. CC and FC performed the cell cultures. CC, TL, KR and NR performed behavioral tests and analyses. CC, FC, SS, NR, LH performed the histology, microscopy and image analyses. CC analyzed Mass Spectrometry and immunoprecipitation data. KR performed viral injection surgery and CC performed spines analysis. MVA performed the analyses of inflammation. FP and CC performed the microdialysis under the supervision of MK. AS and RB performed single-cell RNA sequencing experiments and analysis. CC and NT wrote the manuscript with input from all authors. NT supervised the work.

## Competing interests

The authors have filed a patent application

