## Supplementary Figures for "Astrocyte-derived PEA116 increases adult hippocampal neurogenesis and confers stress resilience"

Supp. Fig. 1

A

| Name | Sequence | MW (Da) | Origin protein | Gene symbol |
|---|---|---|---|---|
| P1 | DGEVIKDSKQEHKDVV | 1825.99 | Glial fibrillary acidic protein | GFAP |
| P2 | KTVETRDGQVINETSQ | 1803.92 | Vimentin | VIM |
| P3 | DPIIEERHGGYQP | 1510.73 | Creatine kinase B | CKB |
| P4 | EDLEQLKSA | 1032.52 | Phosphoprotein enriched in astrocytes | PEA-15 |
| P5 (PEA116) | SEEEIIKLAPPP | 1321.74 | Phosphoprotein enriched in astrocytes | PEA-15 |

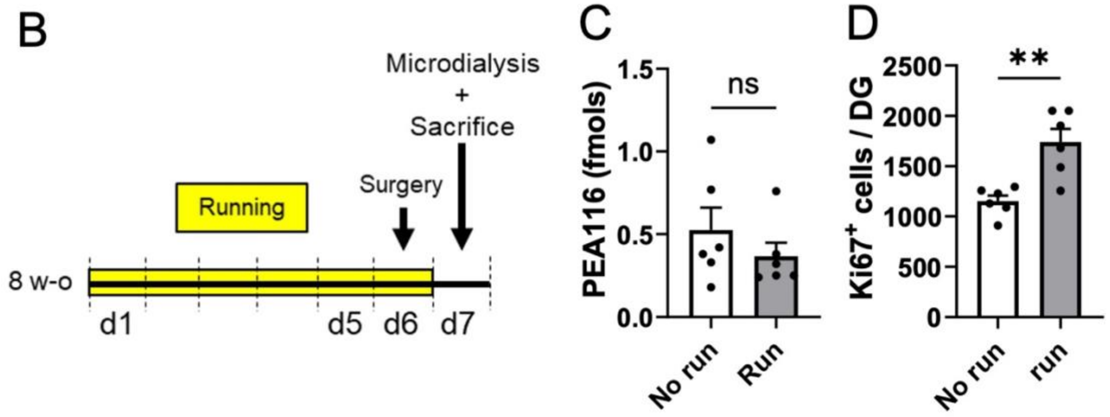


**Supplemental Figure 1: PEA116 is derived from PEA-15 and is found in the extracellular space of the DG. A.** Table of the peptides (sequence, molecular weight, protein of origin) found in ACS by Mass Spectrometry that were redundant across 3 experiments. **B**. Experimental design of the study of the effect of running on the production of the PEA116 peptide *in vivo* (n = 6 mice per group). **C**. Quantification of the PEA116 peptide in microdialysates of the DG in runner and sedentary mice (t = 1.004, p = 0.3392, unpaired t-test, two-tailed, n = 6 mice per group). **D**. Number of Ki67$^{+}$ cells in the DG (t = 4.081, p = 0.0022, unpaired t-test, two-tailed, n = 6 mice per group). Histograms show average ± SEM. ns.: not significant p>0.05; *p<0.05; **p<0.01; ***p<0.001.

Supp. Fig. 2

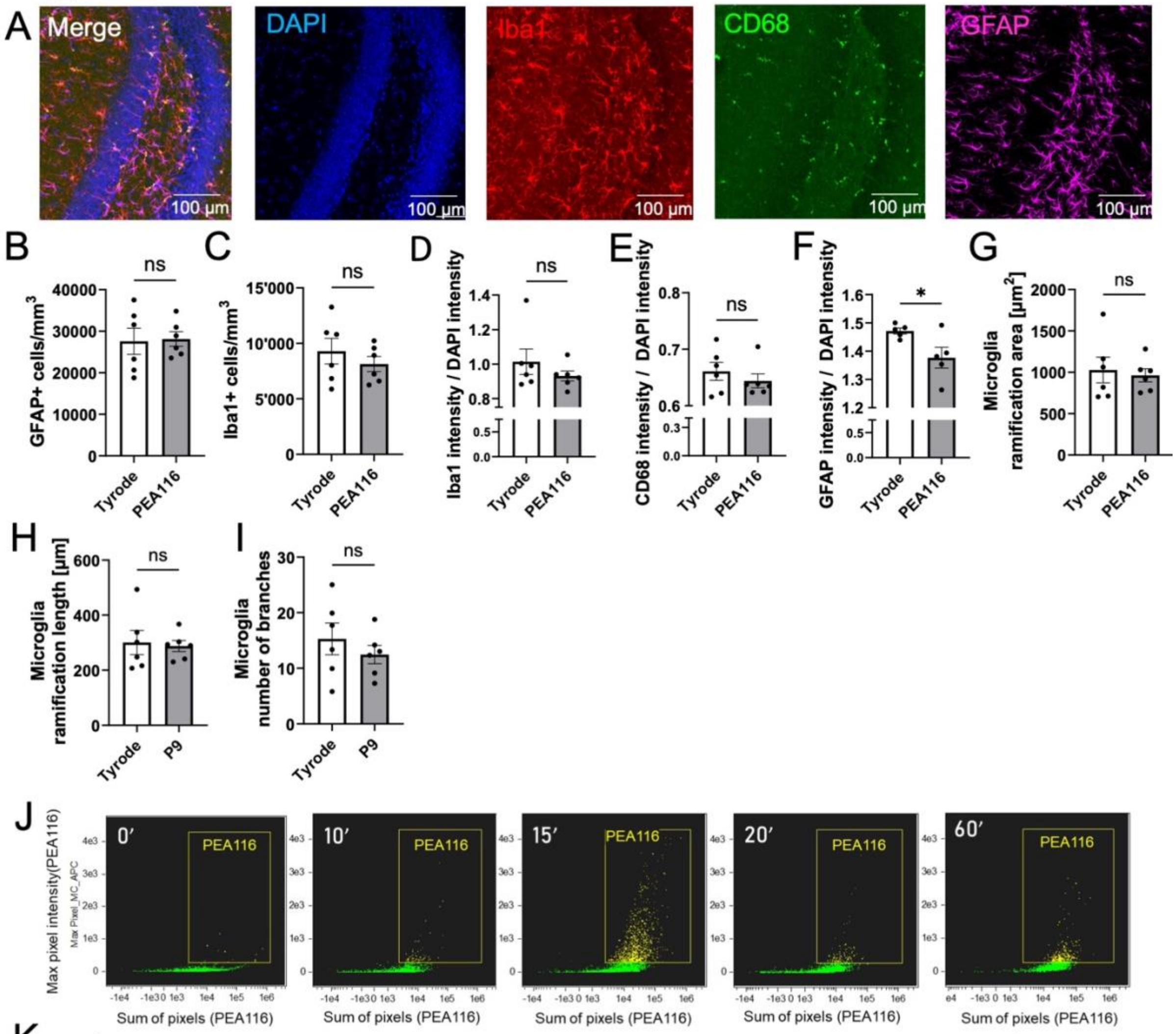


K

| Proteins involved in adult neurogenesis | Protein involved in quiescence | Cell division | Cell cycle | Positive regulation of translation |
|---|---|---|---|---|
| Cdk4 | Huwe1 | Cdk1 | Appl2 | Abcf1 |
| Cdk5 | Ilk | Cdk4 | Cdk5 | Ago2 |
| Dbn1 | | Cdk5 | Csnk1a1 | Cdk4 |
| Dnmt1 | | Ckap5 | Csnk2a2 | Cirbp |
| Fgf2 | | Clasp2 | Gnai2 | Ddx3x |
| Fmr1 | | Csnk1a1 | Gnai3 | Eif4a3 |
| Gsk3b | | Gnai2 | Khdrbs1 | Fmr1 |
| Lgals1 | | Gnai3 | Pelo | Fxr1 |
| Nampt | | Mad1l1 | Ppp1cc | Larp1 |
| Nes | | Mcmbp | Ppm1g | Mapk1 |
| Rac1 | | Numa1 | Rala | Rbm4 |
| Rdx | | Pelo | Rif1 | Ssb |
| | | Ppp1cc | Vrk1 | |
| | | Rala | | |
| | | Rcc1 | | |
| | | Spc25 | | |
| | | Tpx2 | | |
| | | Vrk1 | | |

**Supplemental Figure 2: Effect of PEA116 on Inflammation and immunoprecipitation experiments**. **A.** Confocal micrographs of hippocampal slices immunostained for microglia (Iba1 and CD68) and astrocytes (GFAP) **B-I. B.** Density of astrocytes in the DG of mice treated with Tyrode or PEA116 ($t = 0.1508$, $p = 0.8831$, unpaired t-test, two-tailed, $n = 6$ mice per group). **C.** Density of microglia ($t = 0.8649$, $p = 0.4074$, unpaired t-test, two-tailed, $n = 6$ mice per group). **D-E-F.** Intensity of Iba1, CD68 and GFAP immunostaining as compared to DAPI (**D.** $t = 1.043$, $p = 0.3213$, unpaired t-test, two-tailed, $n = 6$ mice per group; **E.** $t = 0.8481$, $p = 0.4162$, unpaired t-test, two-tailed, $n = 6$ mice per group; **F.** $t = 2.474$, $p = 0.0384$, unpaired t-test, two-tailed, $n = 5$ mice per group). **G.** Microglia ramifications ($t = 0.3692$, $p = 0.7197$, unpaired t-test, two-tailed, $n = 6$ mice per group). **H.** Microglia ramification length ($t = 0.2727$, $p = 0.7906$, unpaired t-test, two-tailed, $n = 6$ mice per group). **I.** Microglia number of branches ($t = 0.8591$, $p = 0.4104$, unpaired t-test, two-tailed, $n = 6$ mice per group). **J.** Flow cytometry images of the detection of the PEA116-HA$^{+}$ aNSPC at different time points after PEA116-HA treatment. **K.** Table of the lists of proteins detected with immunoprecipitation assay and known to be involved in adult neurogenesis according to the MANGO data base, in quiescence, cell division, cell cycle and translation. Histograms show average ± SEM. ns.: not significant $p>0.05$; $*p<0.05$; $**p<0.01$; $***p<0.001$.
